# The Impact of Mismatches Within the RNA: DNA Hybrid in the Transposon-Encoded Type I-F CRISPR-Cas System

**DOI:** 10.1101/2024.02.01.578492

**Authors:** Amnah Alalmaie, Raed Khashan

## Abstract

Recent research has revealed a collaboration between type I-F CRISPR systems and Tn7 transposons in certain bacteria, leading to the discovery of a new gene-editing tool called INTEGRATE. This system integrates transposons into the target strand without introducing a double-strand break or repair mechanism, making it highly promising. The published results showed that a full match between the spacer region of crRNA and the target DNA is necessary to form a stable and complete R-loop, which is critical for accurate integration. PAM-distal mismatches affected RNA-guided DNA transposition differently, with mismatches in positions 25-28 completely blocking the process, while mismatches in positions 29-32 were tolerated. To understand the impact of PAM-distal mismatches on the R-loop stabilization and transposition process, classical all-atom molecular dynamics simulations were conducted using three independent models, including one with a complete RNA-DNA match and two with PAM-distal mismatches located at positions 25-32. Our results suggest that the stable rotation of Cas8-HB is a critical step in the interaction with TniQ and initiation of the DNA transposition process. Network analysis techniques were employed to investigate the communication pathways within Cas8 and TniQ dimers, including eigenvector centrality and correlation analysis. These techniques revealed that certain amino acids in TniQ and Cas8 were highly central to the communication pathways, as they exhibited significant changes in centrality under both mismatches. The findings discussed in this study provide valuable insights into the mechanisms involved in the DNA transposition process and shed light on how PAM-distal mismatches can affect these mechanisms.

## I. Introduction

The CRISPR system, short for “Clusters of Regularly Interspaced Short Palindromic Repeats,” is an integral part of the bacterial defense mechanism against phage invasion [1,2]. The CRISPR array consists of conserved, palindromic genetic sequences known as “repeats” that are interspersed with unique “spacer” sequences [2]. The CRISPR-Cas system is divided into two main classes: Class 1 (type I, type III, and type IV) and Class 2 (type II, type V, and type VI) [3]. Class 1 systems have a multi-subunit effector complex composed of several Cas proteins that assemble around mature CRISPR-RNA (crRNA) to form a large protein complex [3]. In contrast, class 2 systems have single nucleases, either Cas9 or Cas12 [3].

While the CRISPR-Cas9 system is a highly efficient gene editing tool, it has a significant limitation - the off-target effect - in which crRNA may guide Cas9 to cut the wrong DNA sequence [4]. The double-strand break in the target DNA caused by Cas9 can be repaired by natural repair mechanisms such as error-prone non-homologous end-joining and homology-directed repair [5]. Many improvements have been made to the CRISPR-Cas system to control the repair process caused by DNA cleavage and to improve the accuracy of CRISPR-Cas tools.

In 2017, a study by Peters et al. revealed the presence of the CRISPR-Cas system in various bacterial TN7 transposons, such as the TN6677 transposon in Vibrio Cholera [6]. This transposon encodes five transposition proteins (TnsA, TnsB, TnsC, and TniQ dimer) and a short CRISPR array similar to the one found in the type I-F variant CRISPR-Cas system [7]. Unlike other CRISPR-Cas systems, the type I-F variant lacks Cas1, Cas2, and Cas3, which are required for spacer acquisition and target cleavage, but it has multi-Cas enzymes known as the Cascade complex (Cas6, six subunits of Cas7, and a fused Cas8/5) [7]. This complex is called the INTEGRATE system and provides a new generation of gene-editing tools that allows for the insertion of large DNA into a genome without causing a double-strand break and not depending on DNA repair mechanisms [7]. Recent successful trials on the INTEGRATE system have demonstrated an impressive on-target transposition accuracy of 95-99% [7].

The RNA-guided DNA transposition mechanism involves four major steps. The first step includes the maturation and processing of crRNA by Cas6, followed by the assembly of the Cascade complex around a 60-nucleotide-crRNA [7–11]. The second step of the RNA-guided DNA transposition mechanism involves the Cascade-crRNA complex binding to the TniQ dimer in a “G” shape, which is twisted helically**, as shown in Fig. 1** [7–11]. One of the distinguishing features of CRISPR-Cas type I is the unique architecture of the Cas7 backbone, where the “thumb” of each Cas7 subunit folds over the crRNA to form a kink in it. This pattern results in a periodic “5+1” pattern, where the sixth nucleotide is flipped out to the side after every five nucleotides [7–12].

**Fig. 1:**
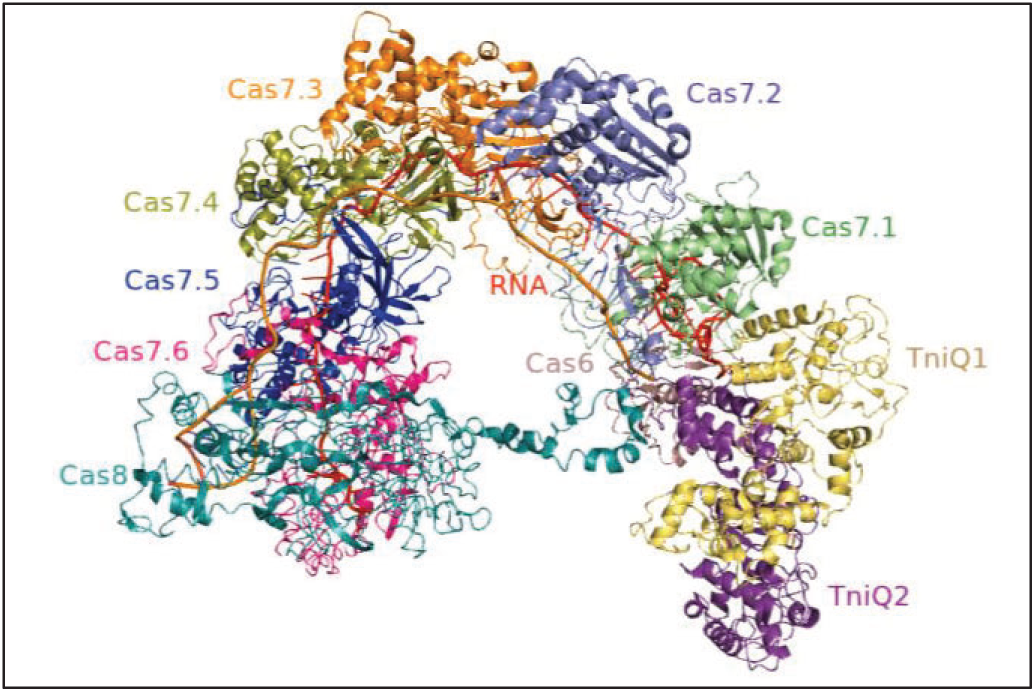
Vc-Cascade-TniQ structure, which adopts a helical (G) architecture where Cas6 forms the head of the complex, six Cas7 subunits form the backbone of the complex, Cas8 forms the tail of the complex in addition to the TniQ dimer. Each monomer of TniQ interacts with the head of the Cascade mainly via Cas6 and Cas7.1 to form a head-to-tail TniQ dimer [12]. Created with PyMol software.

Following the binding of the Cascade-crRNA complex to the TniQ dimer, Cas8 begins scanning the target DNA to locate the Protospacer Adjacent Motif (PAM) sequence. Once the PAM sequence is identified, the spacer region of the crRNA binds to the complementary protospacer segment of the DNA strand, displacing the non-target strand and forming and forming the R-loop Structure [7,9–11,13]. The complete base pairing along the crRNA sequence involves a structural rearrangement of the cascade complex to stabilize the R-loop “R-loop locking,” which will eventually trigger the TniQ activation [14,15]. In the last step, the active TniQ recruits the remaining transposition proteins (TnsC, TnsA, and TnsB). These transposition proteins catalyze the strand cleavage and joining reactions required for transposition, resulting in transposon insertion at the target site [7,9–12]. The TniQ protein is a homodimer located in the Cas operon in the INTEGRATE system, and it works in conjunction with Cas proteins to facilitate precise integration into the intended location [7,12,13]. Each TniQ monomer consists of two significant domains, the N-terminal and the C-terminal. The former is made up of small alpha-helices containing a helix-turn-helix (HTH-1) domain, followed by three beta-sheets that run antiparallel to each other [12,13]. The C-terminal domain is made up of short alpha-helices of varying lengths, a helical domain (HD), and a second helix-turn-helix domain (HTH-2) [12,13].

As mentioned previously, the formation of a complete R-loop serves as a checkpoint to ensure that DNA insertion occurs accurately at the designated “on-target” site where there is complete complementarity between the crRNA and DNA [5,6]. In the INTEGRATE system, the sensitivity of RNA-guided DNA transposition to RNA–DNA mismatches at different regions has been tested experimentally [7] **as shown in Fig. 2**

**Fig 2.**
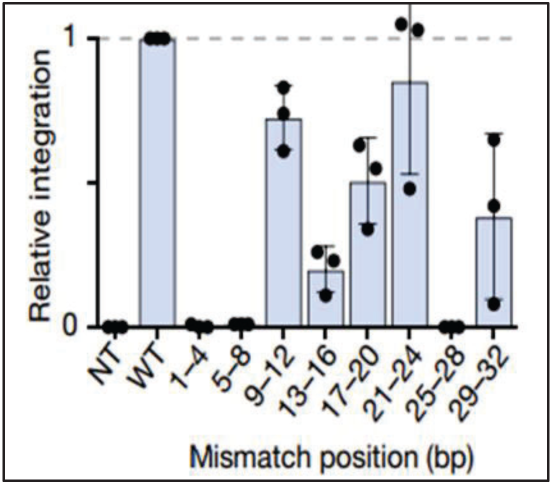
The impact of DNA-RNA mismatches within different regions in the INTEGRATE system. [7] The INTEGRATE system is extremely sensitive to mismatches within the seed sequence, which is located in the PAM-proximal region of the crRNA (specifically positions 1-8 of the spacer region of the crRNA), **as shown in Fig. 3**

**Fig 3.**
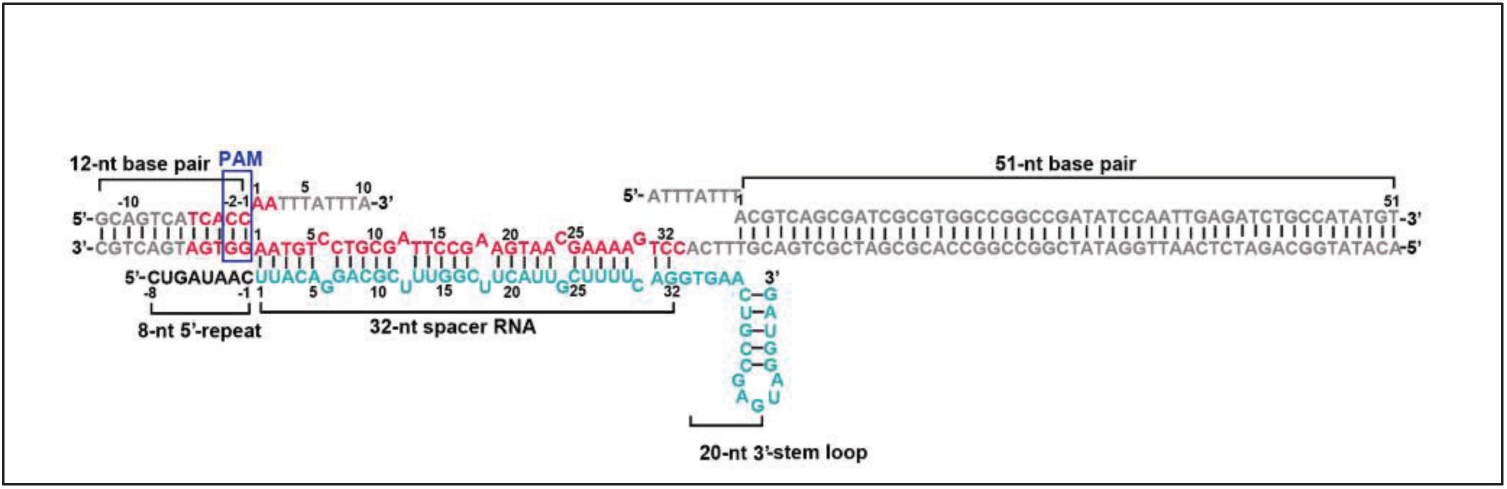
Schematic drawing of the sequences of crRNA and the target DNA [11].

Mismatches within the PAM-proximal region of the crRNA (positions 1-8 of the spacer region) significantly reduce transposition efficiency in the INTEGRATE system, emphasizing the importance of perfect complementarity in this region [7,16–18]. These mismatches cause the Cascade-TniQ complex to quickly dissociate from the target DNA, likely because the R-loop fails to extend beyond the region of mismatches [7,18]. In contrast, PAM-distal mismatches are typically tolerated in type I CRISPR-Cas systems, although they may form unstable partial R-loops [18]. The effects of base pair mismatches on PAM distal sites (positions 25-32) in the INTEGRATE system depend on the location of the mismatch. Mismatches within positions 25-28 completely inhibit RNA-guided DNA transposition, indicating the high sensitivity of this region to mismatches. On the other hand, mismatches within positions 29-32 are tolerated, and the system remains partially functional, with approximately 40% of the transposition efficiency retained [7,12].

To recruit Cas3 nuclease in the type I-F CRISPR-Cas system, Cas8 protein needs to undergo rotation induced by a complete R-loop formation [19]. A similar mechanism is expected in the INTEGRATE system, where the complete R-loop formation triggers the rotation of Cas8, which can activate TniQ. However, TniQ is highly sensitive to mismatches in the PAM distal region (positions 25-28), which could impede the conformational changes necessary for TniQ to recruit TnsC. To better understand the communication pathway and key residues involved in TniQ activation, it is crucial to investigate the allosteric “crosstalk” between Cas8 and TniQ.

Although recent experiments have provided some insights into the effects of RNA-DNA mismatches on the transposition process, the precise conformational changes required for R-loop locking and TniQ activation still need to be fully understood. The stabilization of the R-loop structure and the activation of TniQ may require additional structural rearrangements in the PAM distal region, particularly at positions 25-28. However, determining these conformational changes is a complex experimental task. Molecular Dynamics (MD) simulation provides a promising tool for studying the dynamics of proteins and nucleic acids at the atomic level and could offer valuable insights into the molecular mechanisms underlying the INTEGRATE system. Our study employed all-atom MD simulations to investigate the impact of mismatches in three independent models and provides detailed insight into their effects.

The research investigated the stability and conformational changes of the INTEGRATE system, and explored how mismatches at the PAM and distal regions affect the stability of the RNA-DNA hybrid in the CRISPR-Cas system. Our findings showed that the presence of mismatches can destabilize the RNA-DNA hybrid and interfere with the interactions between the protein subunits and the target DNA strand. The study also found evidence that the stable rotation of Cas8-HB is an essential mechanism in the TniQ interaction and the DNA transposition process. Finally, network analysis techniques helped identify specific amino acid residues that play a crucial role in the communication pathways within Cas8 and TniQ dimers. These findings provide valuable insights into the structural and functional mechanisms involved in the DNA transposition process and how mismatches can affect it. The outcomes of this study are expected to provide valuable guidance to researchers in their efforts to improve the efficacy and precision of the INTEGRATE system.

## II. Simulation Method

Cryo-EM structure is available in the RCSB-PDB database of the ternary TniQ-Cascade complex bound to crRNA and DNA (PDB code:6VBW) [11]. The Structure has an apparent density of 32-nt spacer RNA base-paired to the target strand of DNA [11]. The missing protein residues in the structures were modeled with Charmm-Gui [20].

The introduction of base pair mismatches in the RNA: DNA hybrid has been achieved using the web3DNA software. Specifically, mismatches were created at two positions (25-28) and (29-32) by replacing the ribonucleotides of the PDB ID: 6VBW with non-matching ribonucleotides that do not align with the target DNA [21]. The resultant models have base pair mismatches at position (25-28 or MM11), which are A: DG (at position 25), C: DA (at position 26), C: DA (at position 27), and C: DA (at position 28), whereas the base pair mismatches (29-32 or MM2) are A: DG (at position 29), C: DA (at position 30), C: DA (at position 31), and C: DA (at position 32), **as shown in** **Fig. 4** **Full Match Model (WT)**

**Fig. 4.**
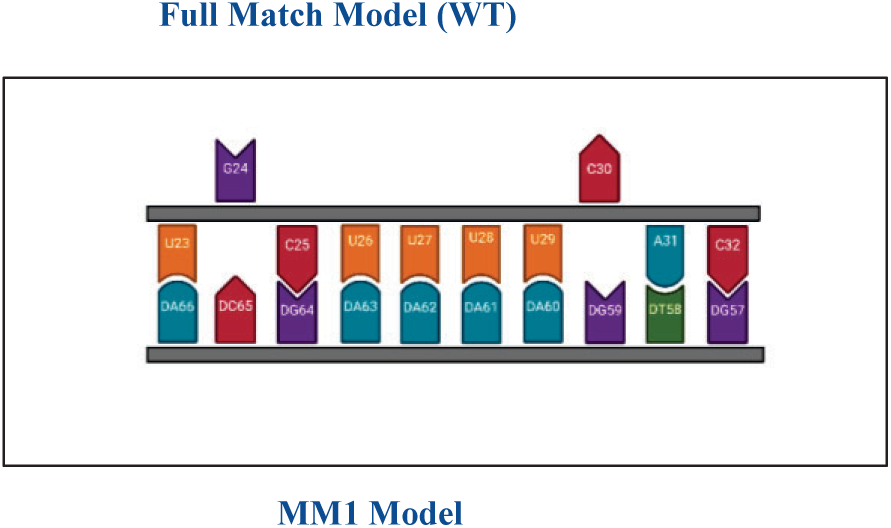
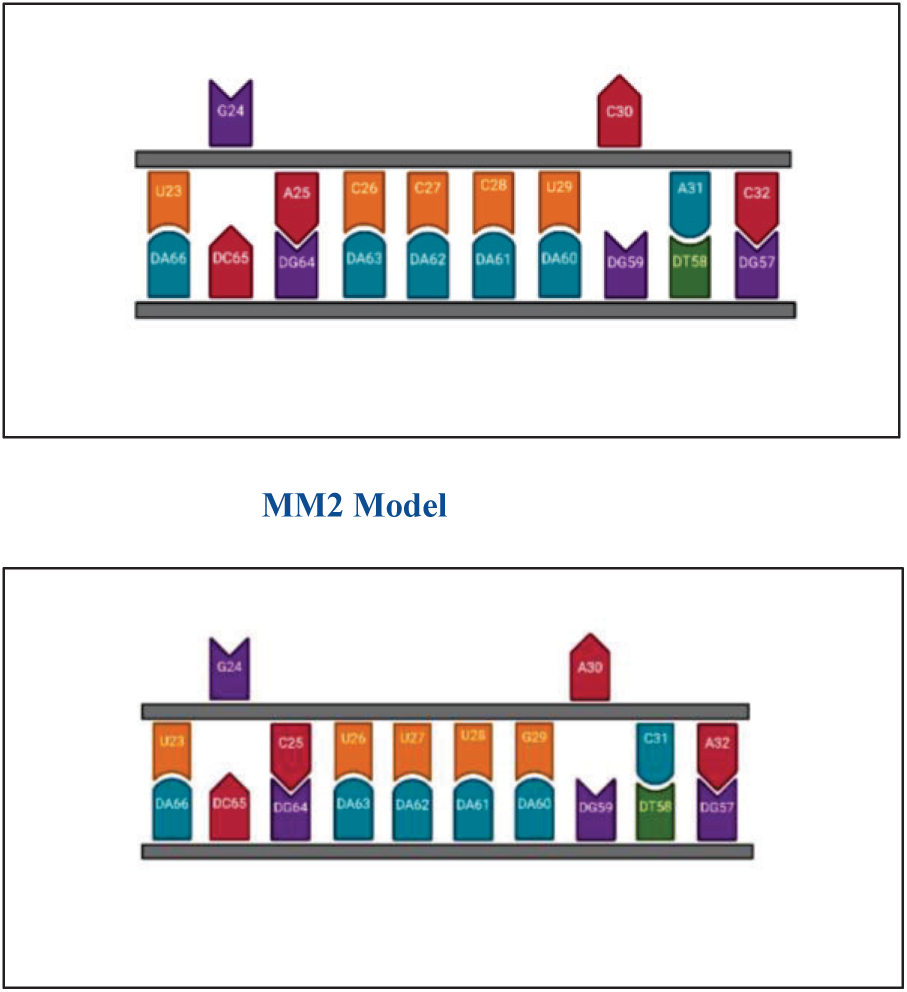
**Schematic representation of the present work** Created with BioRender.com.

The selection of mutations was based on experimental observations. The base pair mismatch at positions 25-28 was chosen as it has been shown to completely inhibit the integration process. On the other hand, the base pair mismatch at positions 29-32 was selected as it still allows for a significant portion (∼40%) of the integration efficiency to be maintained [7].

A popular molecular dynamic package (GROMACS) was used for setting up, running, and analyzing the MD simulations [22]. We conducted three independent simulations for a total of 3 μs. The first simulation represents the fully base-paired state between the spacer region of the crRNA and the target strand of DNA. The second and third simulations, on the other hand, represent the two different mismatch models created as described earlier. All models were prepared and simulated with identical parameters. Amber ff12SB force field was applied for the protein and nucleic acids, including f99bsc0_chiOL3 modification for RNA and ff99bsc0 modification for DNA. This force field has been shown to describe the conformational dynamics of CRISPR MD simulations adequately. The complex was solvated in a triclinic box in each model, and a TIP3P water model was used to simulate water. The simulation box was charge-neutralized by adding Na+ counterions. Each model has ∼ 364453 atoms, including water molecules and ions. Energy minimization was used to remove clashes and bad contacts that would result in an unstable simulation. Fifty thousand (50000) steps of the steepest-descents energy minimization algorithm were used.

Following the energy minimization stage, two equilibrating simulations were run to bring the system temperature and pressure to the desired values. NVT (constant <u>N</u>umber of atoms, <u>V</u>olume, and <u>T</u>emperature) equilibration and NPT (constant <u>N</u>umber of atoms, <u>P</u>ressure, and <u>T</u>emperature) equilibration. In NVT equilibration, the system was heated up from 0 to 300 K in the canonical ensemble (NVT) by running four NVT simulations, **as shown in Table 1**. Temperature control (300 K) was performed by coupling the system to a Nose-Hover thermostat. NPT equilibration was performed for 20 ns. Pressure control was performed by coupling the system to a Parrinello-Rahman barostat at a reference pressure of 1 atm.

**Table 1:** The condition of the four NVT simulations used for the project.

| NVT Simulation Temperature | NVT Simulation Time | Position Restraints |
| --- | --- | --- |
| 0K | 20ps | 300 kcal-mol |
| 0-100K | 50ps | 100 kcal-mol |
| 100-200K | 100ps | 25 kcal-mol |
| 200-300K | 200ps | None (unrestrained) |

An integration time step of 2 fs was used for the production run to conduct MD simulation. For each model, the simulation was carried out in the NPT ensemble for 1 μs of production runs. The Particle-Mesh Ewald (PME) summation method was used to compute the long-range electrostatic interactions, with Fourier spacing of 1.2 nm and fourth-order interpolation. The force-switch method was used to treat the nonbonded interactions (van der Waals) with a cutoff of 1.2 nm. The LINCS algorithm constrained all the hydrogen atoms bonded to the protein-heavy atoms. Our simulations were performed using Shaheen resources. Shaheen is the name of a series of supercomputers owned and operated by the <u>K</u>ing <u>A</u>bdullah <u>U</u>niversity of <u>S</u>cience and <u>T</u>echnology (KAUST) in Saudi Arabia.

## III Results

### Structural Evaluation

To assess the initial stability of the simulations, we calculated the root mean square deviation (RMSD) of the backbone atoms of the entire complex and each component of the system. The RMSD was calculated relative to the first frame of the simulation, which was used as the reference frame, **as shown in Fig. 5**

**Fig. 5:**
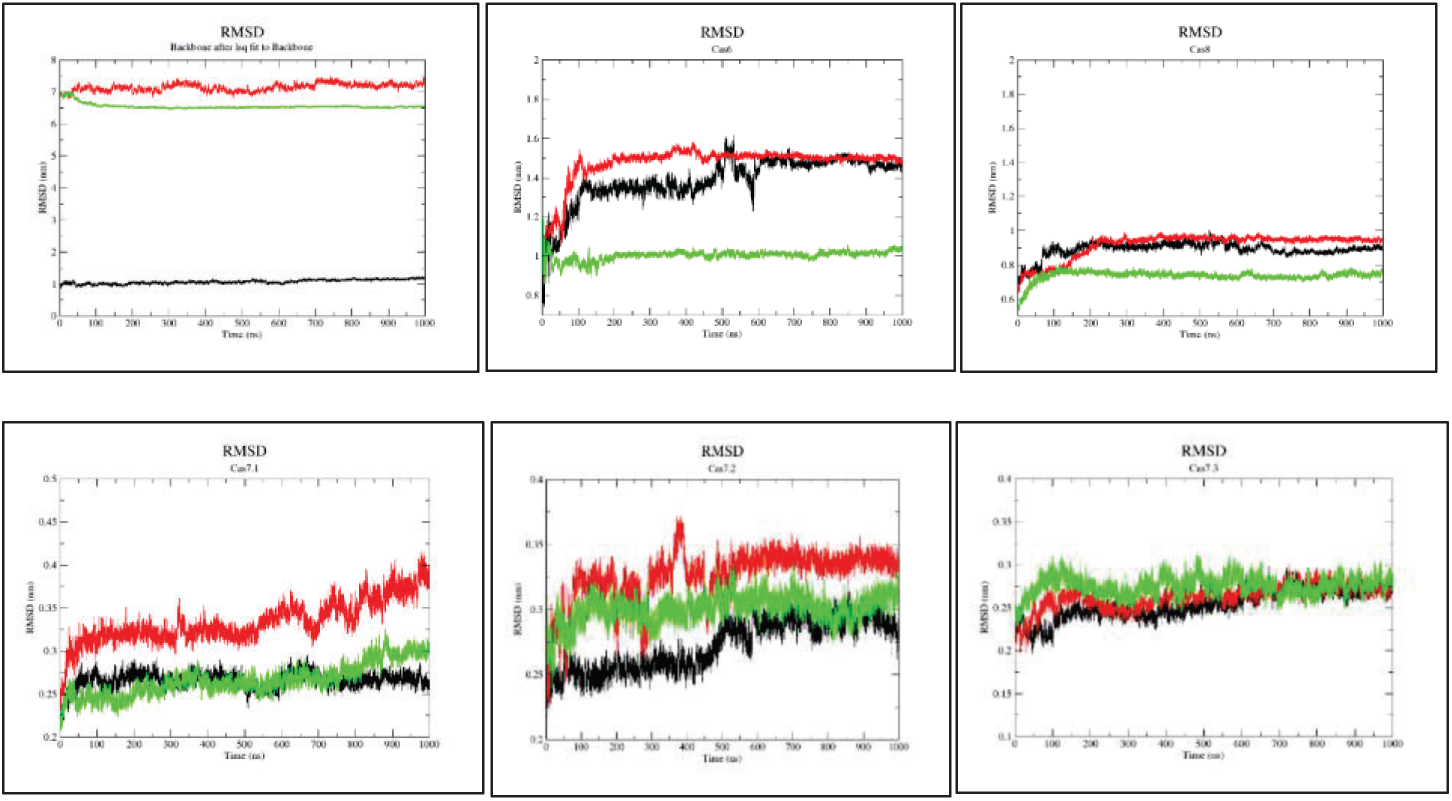
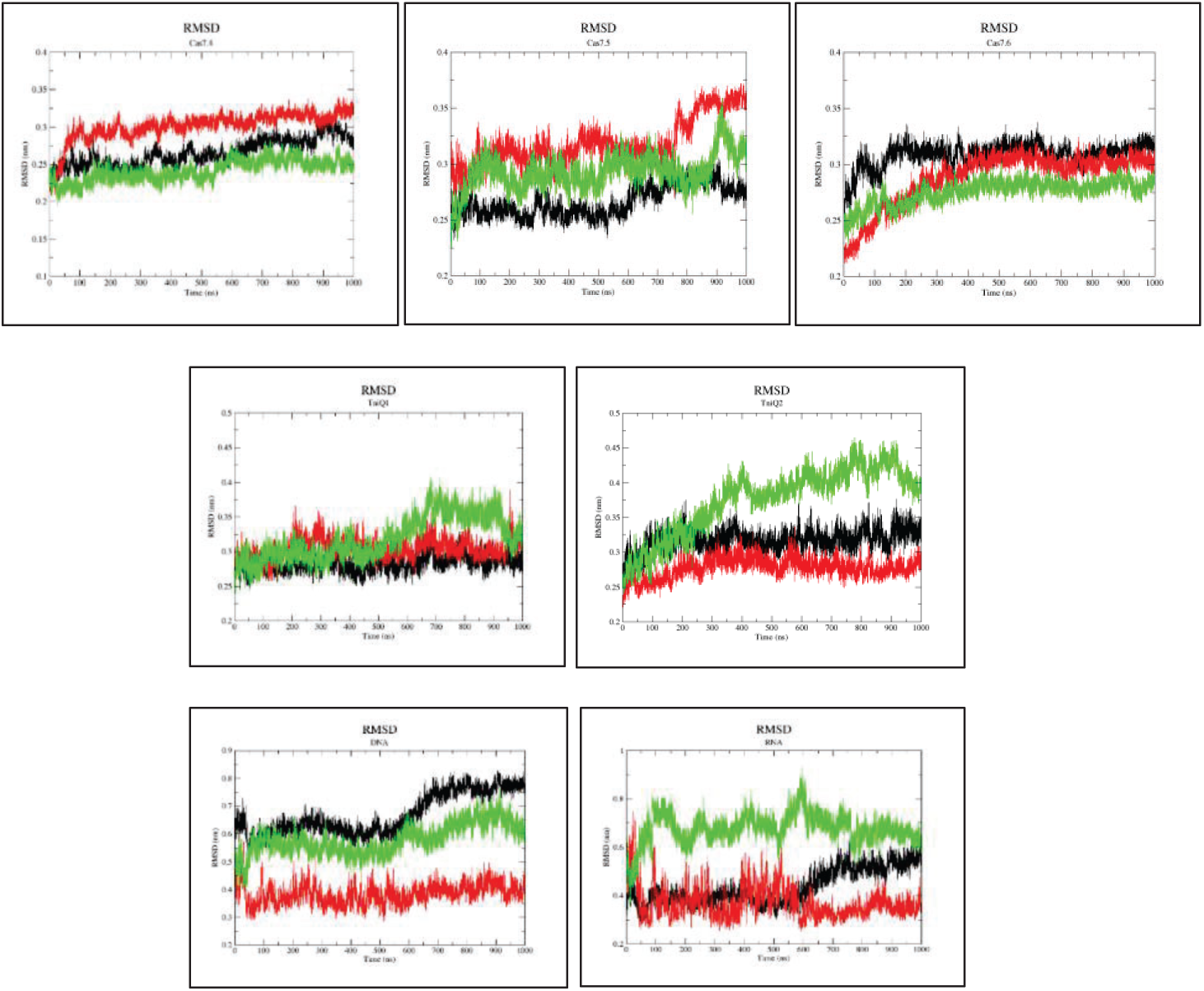
Root-mean-square deviation (RMSD) compared to the first frame over the timescale of 1 μs. Black is the full-match model, red is the MM1 model, and green is the MM2 model.

Throughout the trajectory, the RMSD values of backbone atoms varied to some extent in the MM1 model compared to the other models. Cas6 showed the highest fluctuations in the RMSD, around 500-650 ns, only in the full match model, whereas the RMSD values of Cas8 remained relatively constant in all models. Among the Cas7 subunits, Cas7.1, Cas7.2, and Cas7.5 exhibited the highest fluctuations in their RMSD values. The RMSD values of Cas7.1 remained unstable until 700 ns in the full match and MM2 models but fluctuated in the MM1 model beyond 700 ns. Cas7.2, on the other hand, experienced the highest fluctuations in the RMSD values in the MM1 model. This behavior of Cas7.1 and Cas7.2 is expected, as mismatches were introduced in the RNA-DNA hybrid region near these subunits. The RMSD behavior of Cas7.5 was unexpected, as the RMSD values of this subunit fluctuated in all models. Moreover, the TniQ dimer experienced the most fluctuations in the MM2 model.

To reduce the complexity of the analysis, we have clustered the conformations in the trajectory using an ensemble root mean square deviation (RMSD) clustering algorithm, such as the “*gmx cluster*” algorithm provided by the Gromacs software package. This algorithm groups similar conformations based on a selected RMSD cutoff value, which measures the structural similarity between conformations. After clustering, representative conformations are extracted from each cluster, reducing the conformational size and removing redundant information. The Gromos clustering algorithm is used with different cutoff values to determine the optimal clustering criteria for the specific MD simulation. The clustering was performed for the entire INTEGRATE with different RMSD cutoff values for the last 500 ns of each trajectory, are **shown in Table 2**.

**Table 2:** Effect of the RMSD cutoff values difference on cluster size for the entire INTEGRATE system.

| **RMSD** | **Full Match Model** | **MM1** | **MM2** |
| --- | --- | --- | --- |
| 0.3 nm | 1571 clusters | 17 clusters | 12 clusters |
| 0.4 nm | 1251 clusters | 4 clusters | 5 clusters |
| 0.5 nm | 175 clusters | 3 clusters | 3 clusters |
| 0.6 nm | 19 clusters | 1 cluster | 1 cluster |

The initial clustering analysis showed that the Full match model exhibited more dynamics compared to the mismatch models. The presence of base pair mismatches at PAM distal ends severely impacted the entire complex’s conformational dynamics. This is likely reflected in the clustering analysis, where the number of clusters is reduced for the mismatch models compared to the Full match model.

To gain further insights, we have conducted additional clustering analysis on certain pairs of the INTEGRATE components with an RMSD cutoff of 0.3 nm. This choice of cutoff value implies that conformations with an RMSD value less than 0.3 nm were considered similar and grouped together in a single cluster. This approach allowed for comparing conformational dynamics for the different pairs. The top clusters, which represent at least 70% of the most frequently visited conformations, were identified for each pair of the INTEGRATE components. The specific details on the number of clusters, average RMSD, and percentage of the top clusters **are shown in Table 3 A and B**.

**Table 3:**
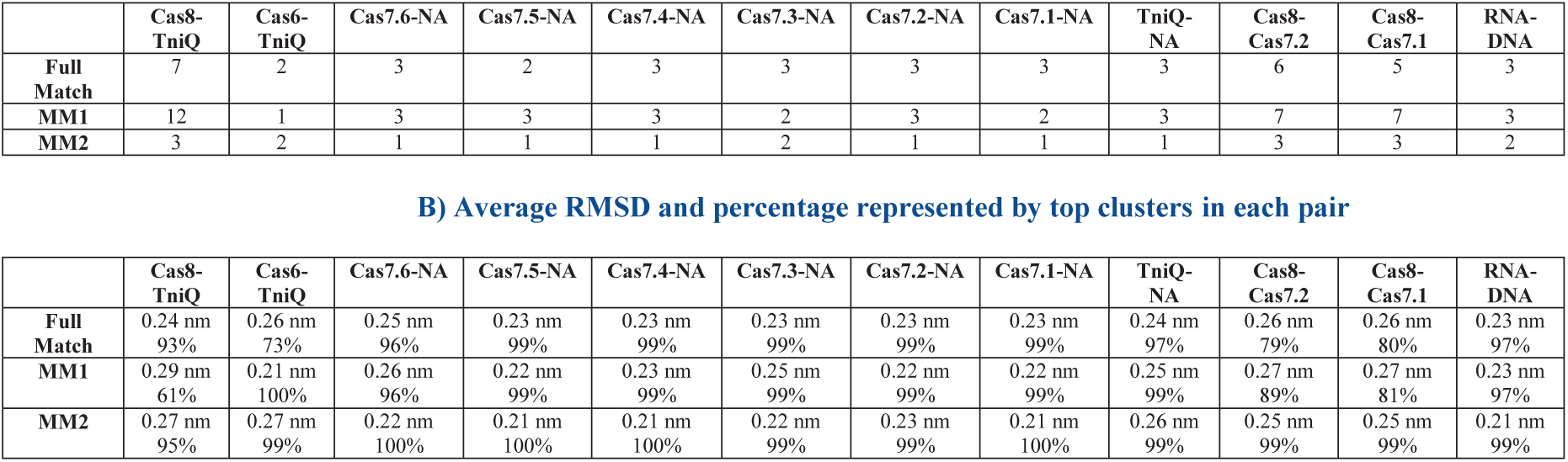
A) Cluster size for different pairs of the INTEGRATE components.

|  | Cas8-TniQ | Cas6-TniQ | Cas7.6-NA | Cas7.5-NA | Cas7.4-NA | Cas7.3-NA | Cas7.2-NA | Cas7.1-NA | TniQ-NA | Cas8-Cas7.2 | Cas8-Cas7.1 | RNA-DNA |
| --- | --- | --- | --- | --- | --- | --- | --- | --- | --- | --- | --- | --- |
| **Full Match** | 7 | 2 | 3 | 2 | 3 | 3 | 3 | 3 | 3 | 6 | 5 | 3 |
| **MM1** | 12 | 1 | 3 | 3 | 3 | 2 | 3 | 2 | 3 | 7 | 7 | 3 |
| **MM2** | 3 | 2 | 1 | 1 | 1 | 2 | 1 | 1 | 1 | 3 | 3 | 2 |

**B) Average RMSD and percentage represented by top clusters in each pair**
|  | Cas8-TniQ | Cas6-TniQ | Cas7.6-NA | Cas7.5-NA | Cas7.4-NA | Cas7.3-NA | Cas7.2-NA | Cas7.1-NA | TniQ-NA | Cas8-Cas7.2 | Cas8-Cas7.1 | RNA-DNA |
| --- | --- | --- | --- | --- | --- | --- | --- | --- | --- | --- | --- | --- |
| **Full Match** | 0.24 nm<br>93% | 0.26 nm<br>73% | 0.25 nm<br>96% | 0.23 nm<br>99% | 0.23 nm<br>99% | 0.23 nm<br>99% | 0.23 nm<br>99% | 0.23 nm<br>99% | 0.24 nm<br>97% | 0.26 nm<br>79% | 0.26 nm<br>80% | 0.23 nm<br>97% |
| **MM1** | 0.29 nm<br>61% | 0.21 nm<br>100% | 0.26 nm<br>96% | 0.22 nm<br>99% | 0.23 nm<br>99% | 0.25 nm<br>99% | 0.22 nm<br>99% | 0.22 nm<br>99% | 0.25 nm<br>99% | 0.27 nm<br>89% | 0.27 nm<br>81% | 0.23 nm<br>97% |
| **MM2** | 0.27 nm<br>95% | 0.27 nm<br>99% | 0.22 nm<br>100% | 0.21 nm<br>100% | 0.21 nm<br>100% | 0.22 nm<br>99% | 0.23 nm<br>99% | 0.21 nm<br>100% | 0.26 nm<br>99% | 0.25 nm<br>99% | 0.25 nm<br>99% | 0.21 nm<br>99% |

### Behavior of the DNA-RNA Hybrid in the Presence of the PAM-distal Mismatch

In the full match model, the Watson-Crick base pairing is maintained along the entire heteroduplex, and the PAM distal region (positions 25-32) is stabilized by Cas7.3, Cas7.2, and Cas7.1, **as shown in Fig.6A**. In the MM1 model, mismatches in positions 25-28, the Watson−Crick base pairing was disrupted except for position 28, in which the Cytosine maintains the base paring with the Adenine located at position 61 of the DNA-target strand, regardless of the mismatch, **as shown in Fig. 6B**. Interestingly, we noticed a loss in the Watson-Crick hydrogen bonding at the base-pair positioned at A31: DT58, near the mismatch site (A31: DT58). We also noticed that the Thymine (DT58) was flipped in the MM1 model, **as shown in Fig.6B and C**. In the MM2 model, mismatches in positions 29-32, the Watson−Crick base pairing was maintained only at the base-pair positioned at G29:DA60. However, no base flipping was observed, **as shown in Fig. 6D**.

**Fig. 6.**
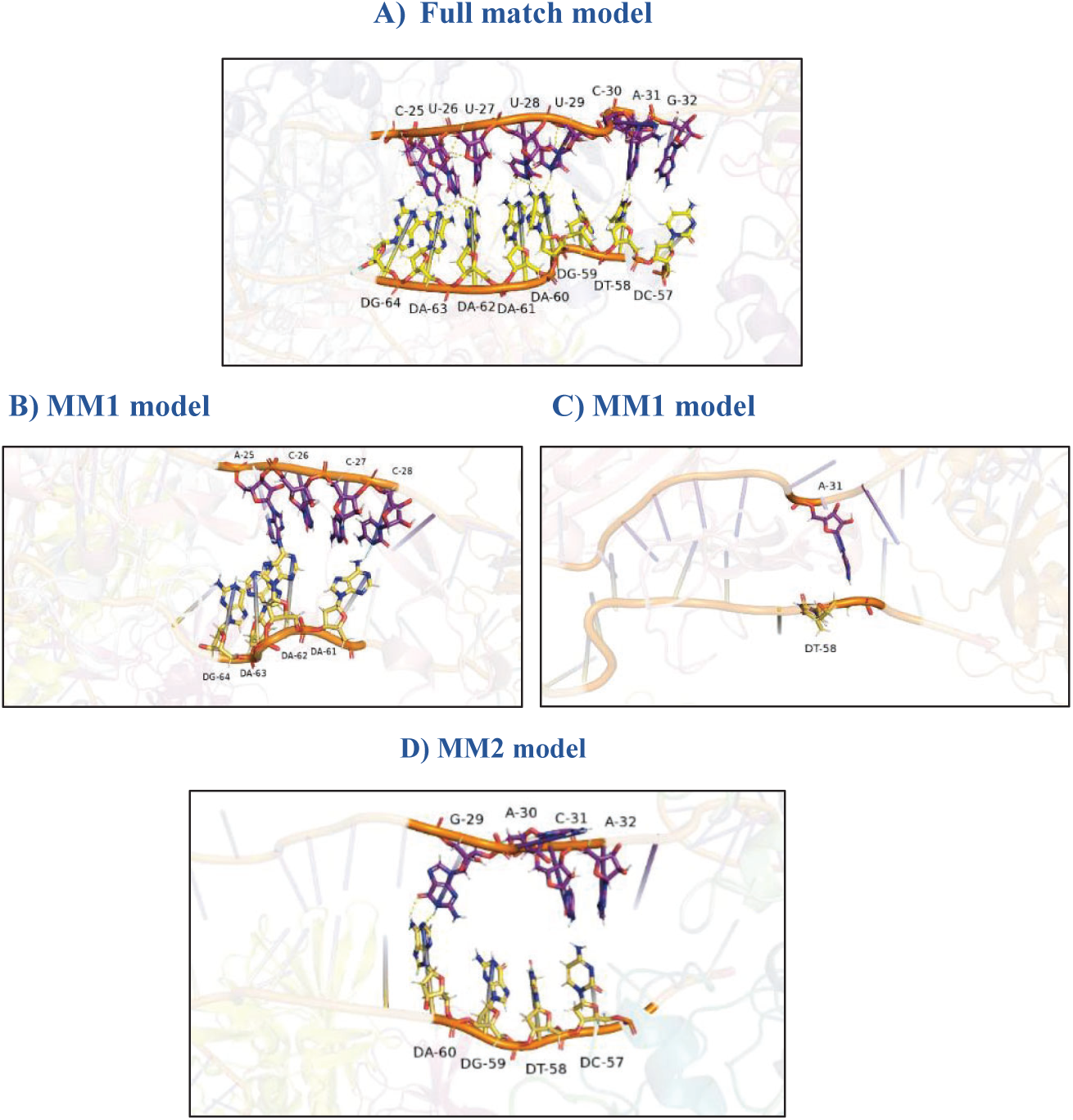
The Behavior of the DNA-RNA base pairing in all models.

The impact of PAM distal mismatches on RNA-DNA hybrid stability was investigated using COCOMAPS, (bioCOmplexes COntact MAPS) a web server used for detailed interaction analysis [23]. The top clusters for three pairs of INTEGRATE components, namely the target strand of the DNA and protein subunits Cas7.3, Cas7.2, and Cas7.1, were analyzed using a cutoff distance of 0.8 nm. This allowed for the identification of both newly formed and disappearing interactions.

Results are shown in **Table 4**.

**Table 4:**
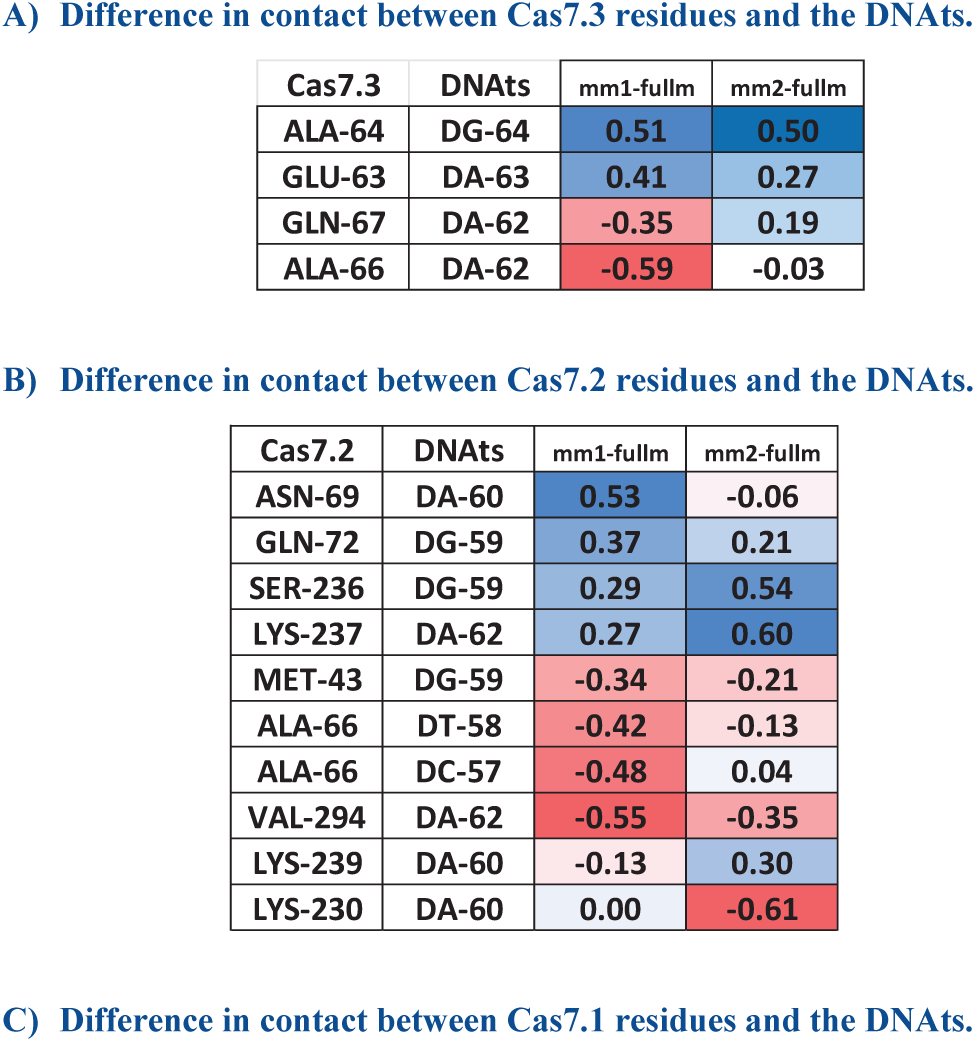
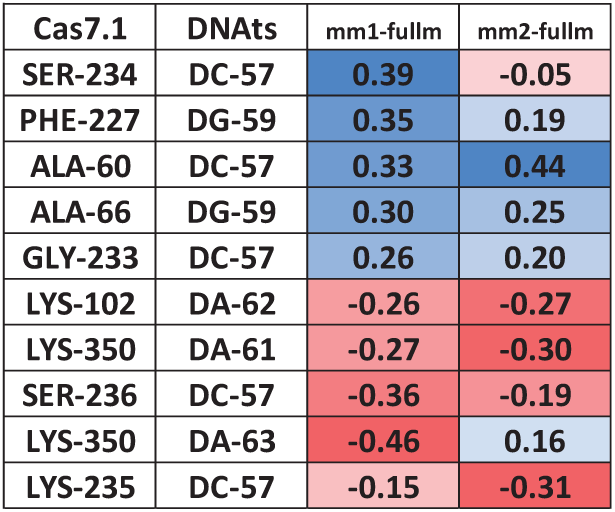
The difference in contacts (in nanometer) between the atoms in each pair was calculated by subtracting the minimum distances in the full match model from those in mismatch models. Only the interactions significantly affected by the PAM distal mismatch are shown. The dark red color indicates newly formed interactions, and the dark blue color indicates disappearing interactions.

Regarding the interaction between Cas7.3 and the target strand, it was found that in the full match model, the interactions between Ala64 and Glu63 of Cas7.3 and the DNA were maintained. However, in the MM1 and MM2 models, the distance between these residues and the DNA increased, indicating a loss of interaction. On the other hand, the distance between Gln67 and Ala66 showed a significant decrease in the MM1 model, suggesting the formation of new interactions between these residues and the DNA.

The interaction between Cas7.2 and the target strand revealed that Asn69 maintained its interaction with the DNA in both the full match and MM2 models but showed a longer distance in the MM1 model, suggesting a possible loss of interaction. On the other hand, Gln72, Ser236, and Lys237 displayed increased distances in the MM1 and MM2 models, indicating a loss of interaction. In contrast, Met43 showed a shorter distance in the MM1 and MM2 models, implying the formation of a new interaction. Furthermore, in the MM1 model, Ala66 displayed a shorter distance and formed new interactions with the Cytosine and Thymine positions at 57 and 58, respectively. Val294 interacted with the Adenine positions at 62 in both MM1 and MM2 models, with shorter distances indicating newly formed interactions. Lys230 displayed a significant shortening in distance to the Adenine position at 60 in the MM2 model, resulting in the formation of a salt bridge.

In the full match model, Lys230 formed a salt bridge with the Guanine position at 64, but this interaction was disrupted in both the MM1 and MM2 models.

When examining the interaction between Cas7.1 and the DNA, it was found that Phe227, Ala66, Ala60, and Gly233 exhibited elongation in the distance, indicating a loss of interaction, in both MM1 and MM2 models. Conversely, Lys102, Lys350, Lys235, Ser236 showed shortening in the distance in both models, suggesting newly formed interactions. Furthermore, Ser234 maintained its interaction with the Cytosine positioned at 57 in the full match and MM2 models, but the interaction was lost in the MM1 model.

It was also observed that the DNA target strand exhibited a “locked” conformation in the full match and MM2 models, wherein Cas7.2 and Cas7.1 fold over the DNA target strand. However, in the MM1 model, the target strand is not locked, and instead, the Guanine positioned at 64 and the

Adenine positioned at 63 were stretched out, as shown in **Fig. 7**. The same phenomenon of target-strand locking was observed in the type I-F and I-E CRISPR-Cas systems [19,24]. The nucleotide stretch observed in the MM1 model is believed to be caused by the absence of a salt bridge between Lys230 and the PAM distal region [25–28]. Salt bridges are electrostatic interactions between acidic and basic groups that play a crucial role in stabilizing protein structures [25]. We believe that the absence of this interaction could destabilize the protein-DNA complex, resulting in the stretched nucleotides observed in the model.

**Fig. 7.**
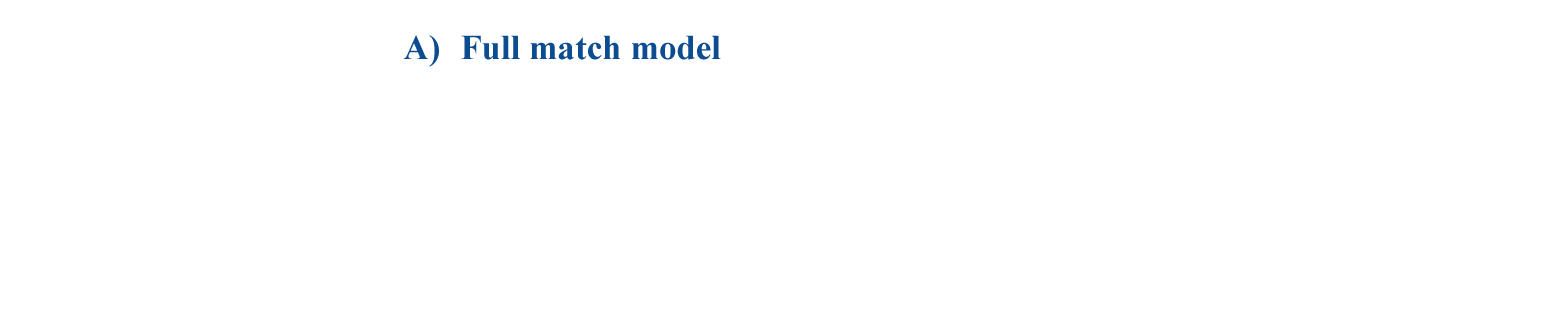
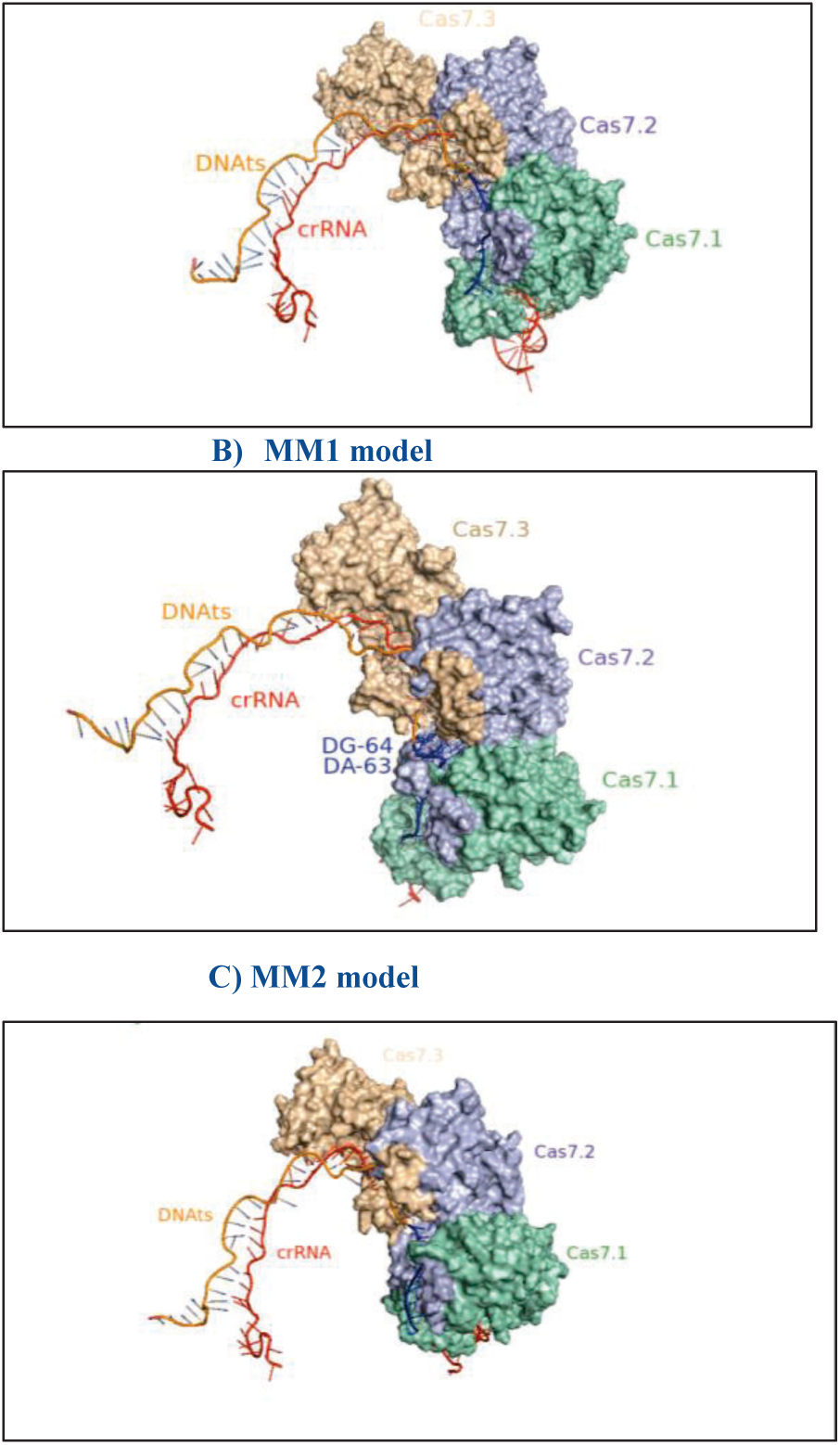
The DNA target strand locks due to the interaction with Cas7.2 and Cas7.1

### Rotation of Cas8-Helical Bundle

Cas8 is composed of three distinct domains: the N-terminal domain (Cas8-NTD, amino acids 1-275), which is responsible for the recognition of the PAM sequence, the middle helical bundle (Cas8-HB, amino acids 276-384), and the C-terminal domain (Cas8-CTD, amino acids 385-640) [9,10]. When the R-loop is completely formed, it is anticipated that the Cas8 helical bundle (Cas8-HB) will rotate and activate TniQ, initiating the DNA transposition process [9,10]. According to the similarities in structure between the INTEGRATE system and type I-F CRISPR-RNA-guided surveillance complex, it is believed that stable rotation of Cas8-HB is important for its interaction with TniQ [19,24]. In the full match and MM2 models, but not the MM1 model, it was observed that the helical bundle of Cas8 underwent a rotation and approached TniQ, resulting in an interaction between the two proteins. Furthermore, the rotated conformation is stabilized by the interaction with Cas7.2. These observations suggest that stable rotation of Cas8-HB is a conserved mechanism that plays a crucial role in the interaction with TniQ and the initiation of the DNA transposition process in both the full match and MM2 models. The COCOMAPS analysis was employed to identify the minimum distance contact between the residues of Cas8-HB and TniQ, using a cutoff distance of 0.4 nm based on the top cluster that represented approximately 70% of the visited conformations [23]. The observed intearctins btween Cas8-HB and the TniQ are shown in Fig. 8

**Fig. 8.**
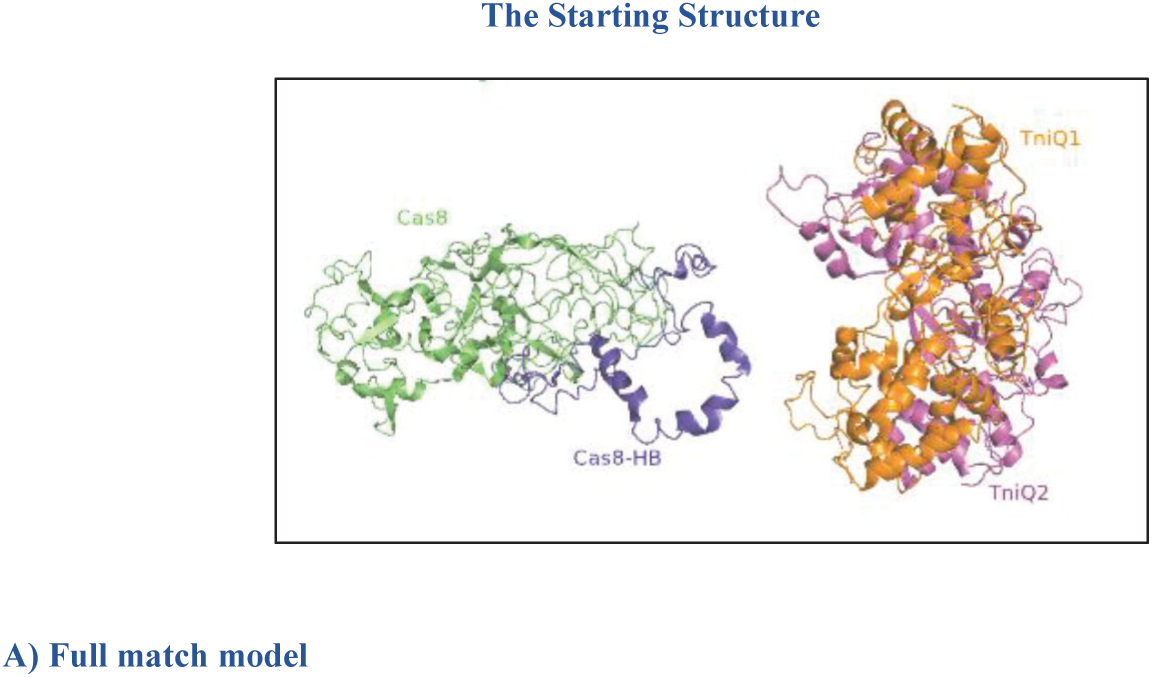
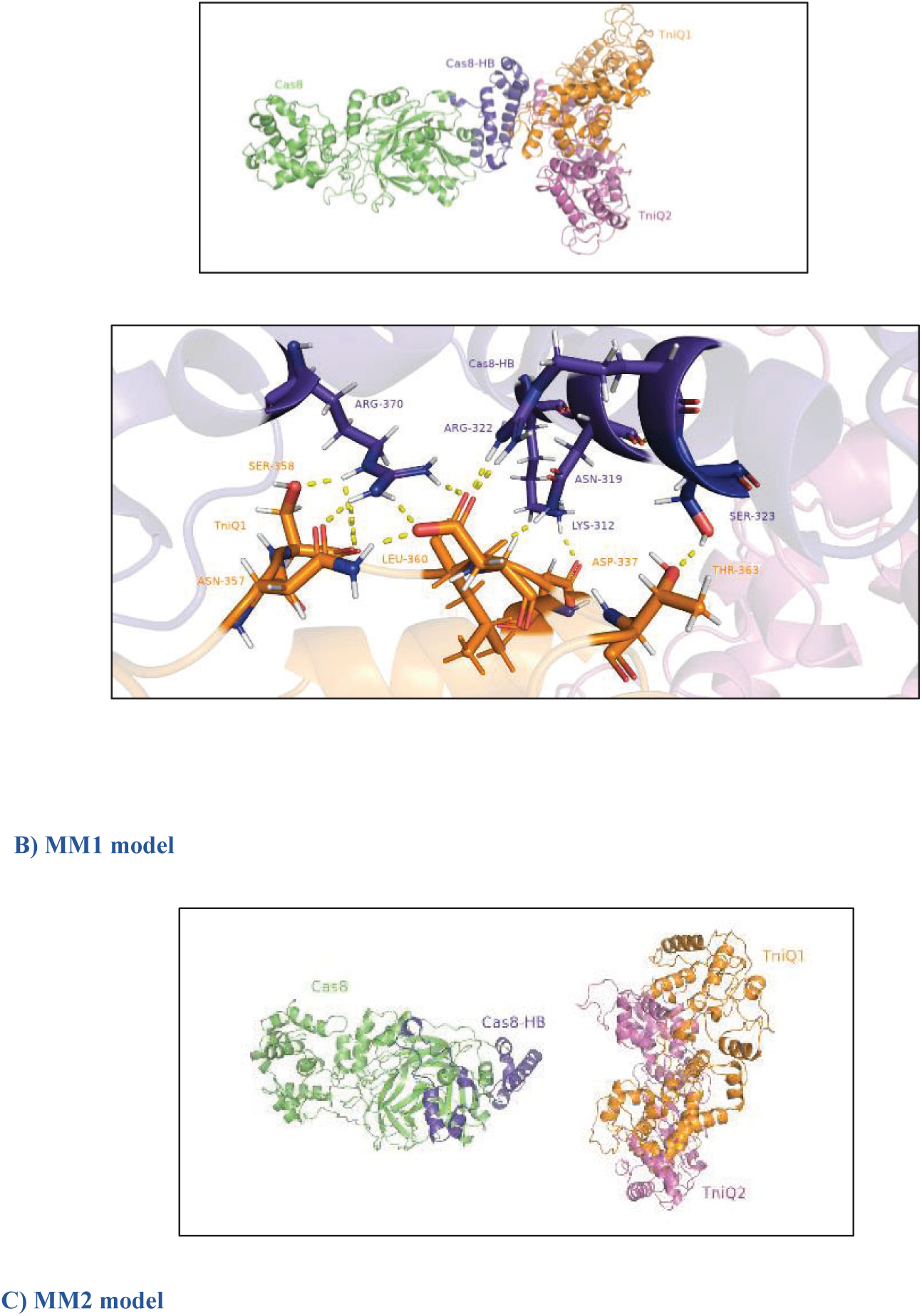
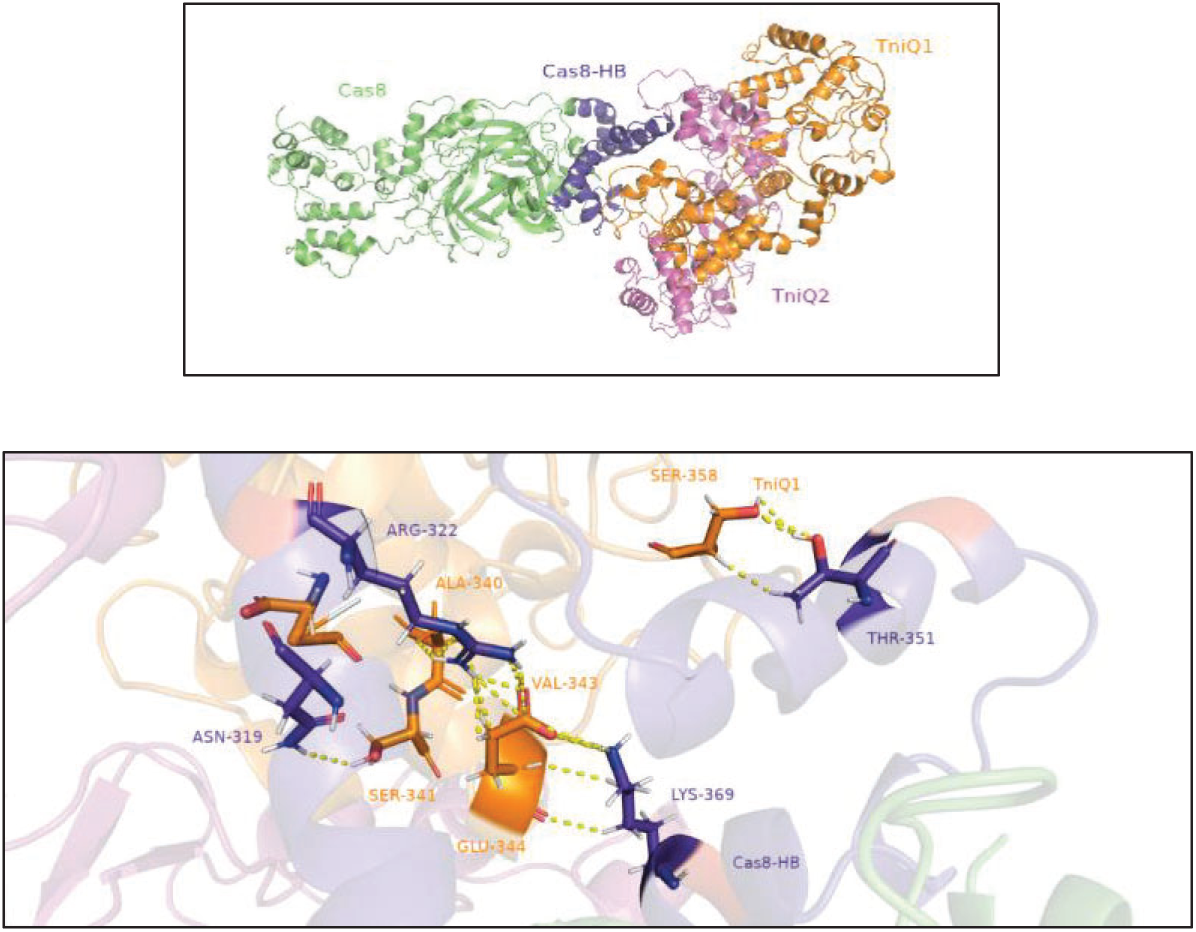
The rotation of Cas8-Helical bundle is stabilized by the interaction with TniQ.

In the full match models, the Cas8-HB residues that predominantly interacted with TniQ1 were Lys312, Asn319, Arg322, Ser323 and Arg370, whereas Arg319 and Arg322, Lys351, and Lys369 being the primary interacting residues of the Cas8-HB in the MM2 model. These residues are part of the helix-turn-helix (HTH) domain of TniQ, which is thought to interact with the Cas8-HB based on the experimental findings. [9].

A COCOMAPS analysis was also performed within a cutoff distance of 0.8 nm. The analysis indicated that Cas8HB and TniQ1 share pair of interactions in both models, while there are no common interactions observed between Cas8HB and TniQ2 in either model, **result is shown in Table 5**.

**Table 5.**
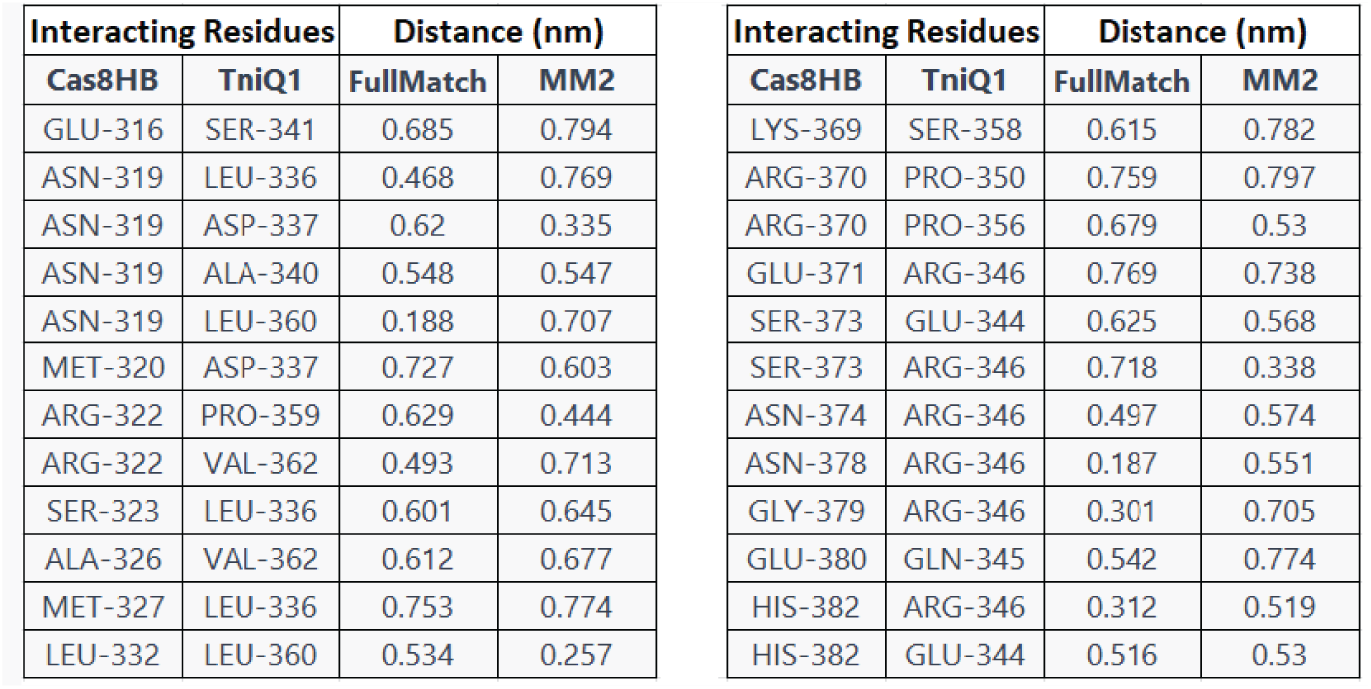
The distance in nm between the Cas8-HB residues that interact with TniQ1, which are common between the Full Match and MM2 models.

| Interacting Residues |  | Distance (nm) |  |
| --- | --- | --- | --- |
| Cas8HB | TniQ1 | FullMatch | MM2 |
| GLU-316 | SER-341 | 0.685 | 0.794 |
| ASN-319 | LEU-336 | 0.468 | 0.769 |
| ASN-319 | ASP-337 | 0.62 | 0.335 |
| ASN-319 | ALA-340 | 0.548 | 0.547 |
| ASN-319 | LEU-360 | 0.188 | 0.707 |
| MET-320 | ASP-337 | 0.727 | 0.603 |
| ARG-322 | PRO-359 | 0.629 | 0.444 |
| ARG-322 | VAL-362 | 0.493 | 0.713 |
| SER-323 | LEU-336 | 0.601 | 0.645 |
| ALA-326 | VAL-362 | 0.612 | 0.677 |
| MET-327 | LEU-336 | 0.753 | 0.774 |
| LEU-332 | LEU-360 | 0.534 | 0.257 |
| Cas8HB | TniQ1 | FullMatch | MM2 |
| LYS-369 | SER-358 | 0.615 | 0.782 |
| ARG-370 | PRO-350 | 0.759 | 0.797 |
| ARG-370 | PRO-356 | 0.679 | 0.53 |
| GLU-371 | ARG-346 | 0.769 | 0.738 |
| SER-373 | GLU-344 | 0.625 | 0.568 |
| SER-373 | ARG-346 | 0.718 | 0.338 |
| ASN-374 | ARG-346 | 0.497 | 0.574 |
| ASN-378 | ARG-346 | 0.187 | 0.551 |
| GLY-379 | ARG-346 | 0.301 | 0.705 |
| GLU-380 | GLN-345 | 0.542 | 0.774 |
| HIS-382 | ARG-346 | 0.312 | 0.519 |
| HIS-382 | GLU-344 | 0.516 | 0.53 |

The rotated conformation of Cas8-HB is stabilized by the interaction with Cas7.2, folding over the DNA target and making contact with the helical bundle of Cas8 to completely lock the target strand as observed in the full match model and MM2 model. The interaction between Cas8-HB and Cas7.2 **are shown in Fig.9**

**Fig. 9.**
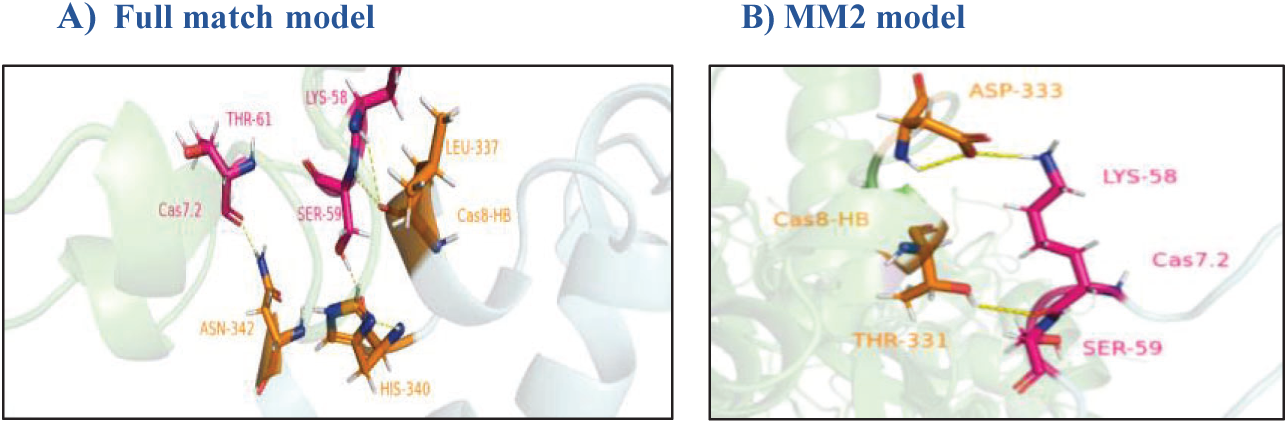
The interaction between Cas8-HB and Cas7.2.

### Revealing the Allosteric Effect of PAM-distal Mismatch on the Dynamics of

To understand how the DNA-RNA hybrid mismatches influence the dynamics of Cas8 and TniQ, the protein correlation network (PCN) in the full match model (Wild Type [WT]) and the mismatch models were computed to identify which specific changes are responsible for altering the overall dynamics of the protein. The PCN is based on network theory, which originated from graph theory. In the graph theory, each residue within a protein is represented as a node in the network, and these nodes are interconnected to other nodes through edges. [26,27] By analyzing the PCN, it is possible to identify key residues and pathways that play a role in the communication between different parts of Cas8 and TniQ [28].

PCN was computed using a new software tool called MDigest, which utilized different correlation analysis techniques [29]. Two of these techniques are the generalized correlation coefficient that is based on linearized mutual information (LMI-based GCC) and the generalized correlation coefficient that is based on the Kabsch-Sander formula (KS-based GCC) [27–30]. The LMI-based GCC method calculates the correlation between the displacements of CA (Cα) atoms in a protein structure. By examining the correlation between Cα displacements, it is possible to identify which regions of the protein move together (i.e., are correlated) or move independently (i.e., are anti-correlated), which provides valuable insights into the protein’s dynamics and allostery [28]. The edges connecting correlated nodes are defined based on the strength of their correlation [27].

On the other hand, the KS-based GCC measures the correlation between electrostatic potential across each frame in the trajectory [29,30]. This is done by using the Kabsch-Sander formula, which calculates the energy between hydrogen bond donor and acceptor groups in each frame of a molecular dynamics simulation [29,30]. These techniques provide unique insights into protein dynamics and allostery, capturing diverse aspects of its dynamics. By employing both methods, it is possible to obtain a more comprehensive understanding of the protein’s behavior. The aim is to investigate how the PAM-distal mismatch could affect the communication pathways within Cas8 and TniQ and potentially alter the dynamics.

Correlation matrices for Cα-displacements from simulated trajectories of Cas8 and TniQ were calculated for the wild-type (WT), MM1, and MM2 variants, as well as their differences. To facilitate a more direct comparison of these matrices, we generated “difference matrices” by subtracting one matrix from another. We then evaluated changes in eigenvector centrality values based on the “difference matrix” between the full match model (WT) and mismatches models, **as shown in** **Fig 10**, **11** **and** **12**.

**Fig.10.**
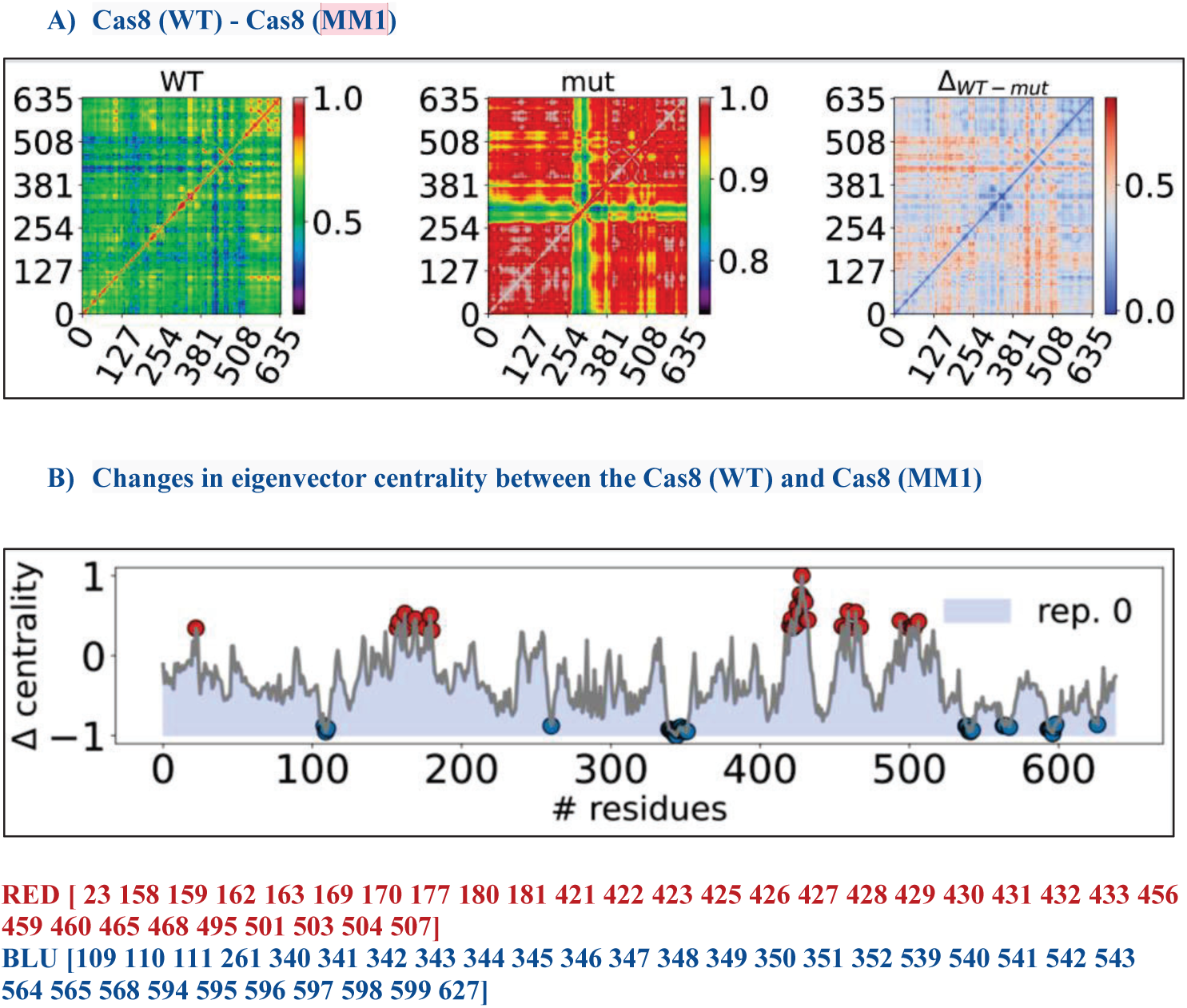
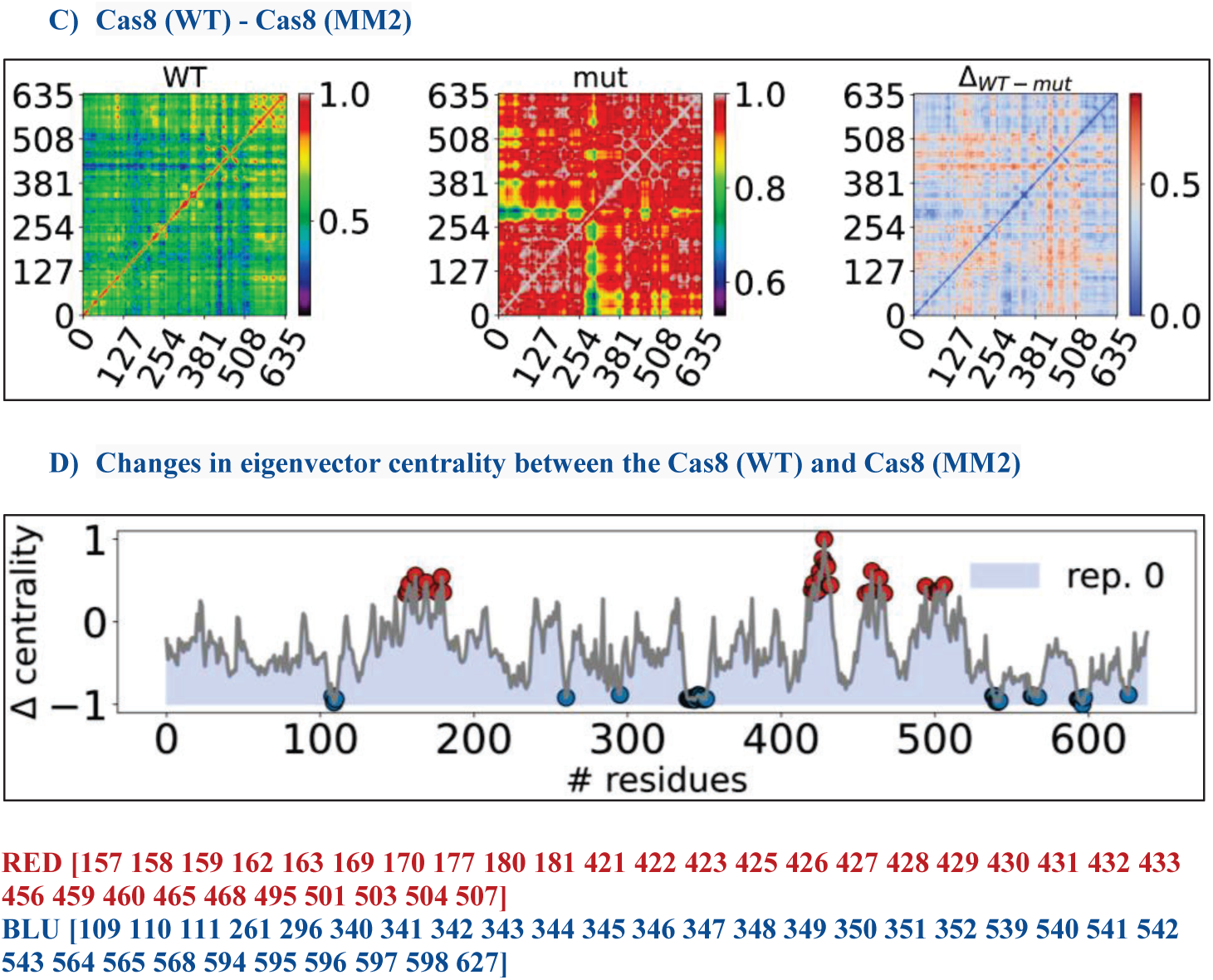
Correlation of Cas8 residues based on LMI metric for the full match model (WT), mismatch models (MM1 and MM2), and their differences.

**Fig.11.**
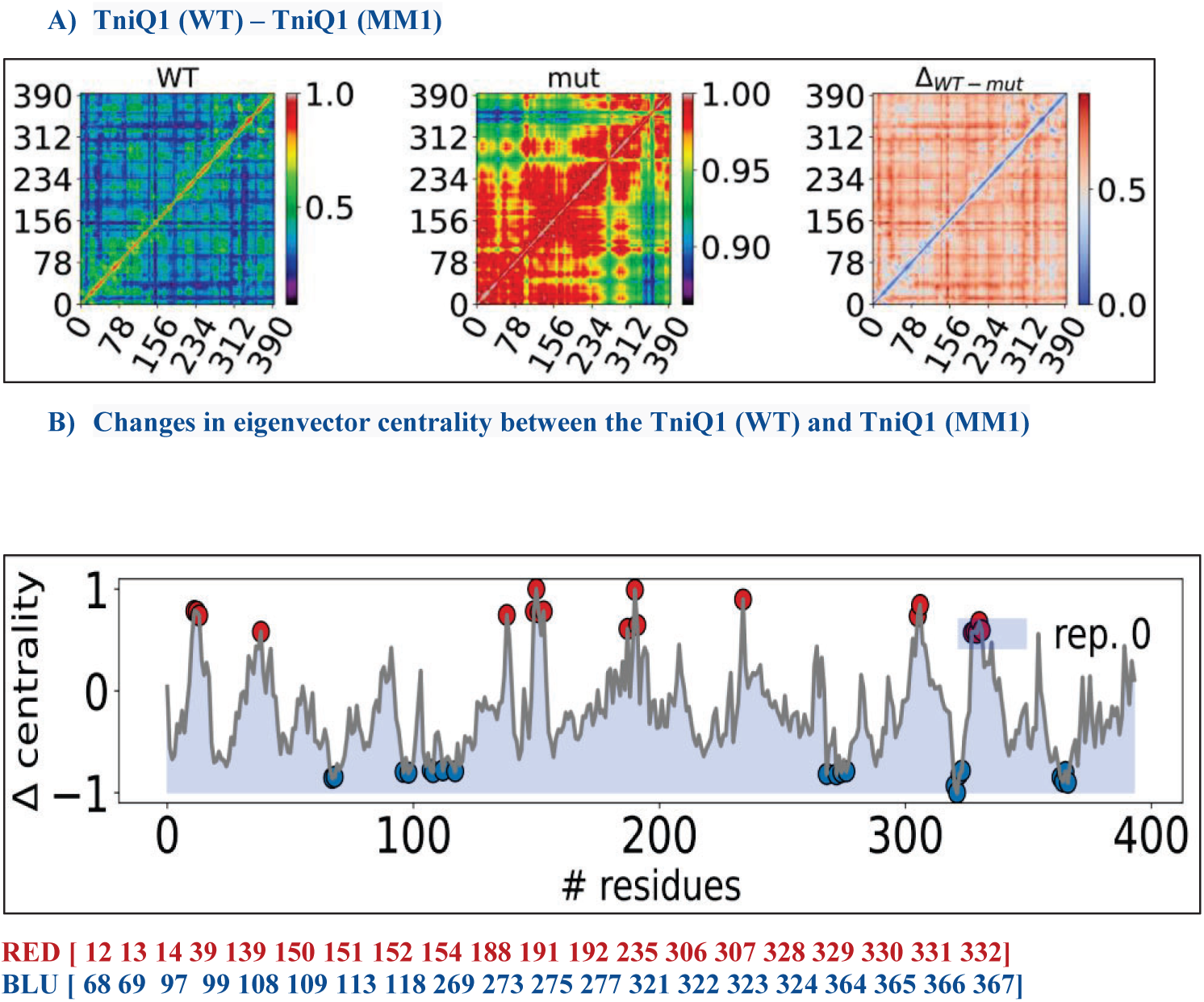
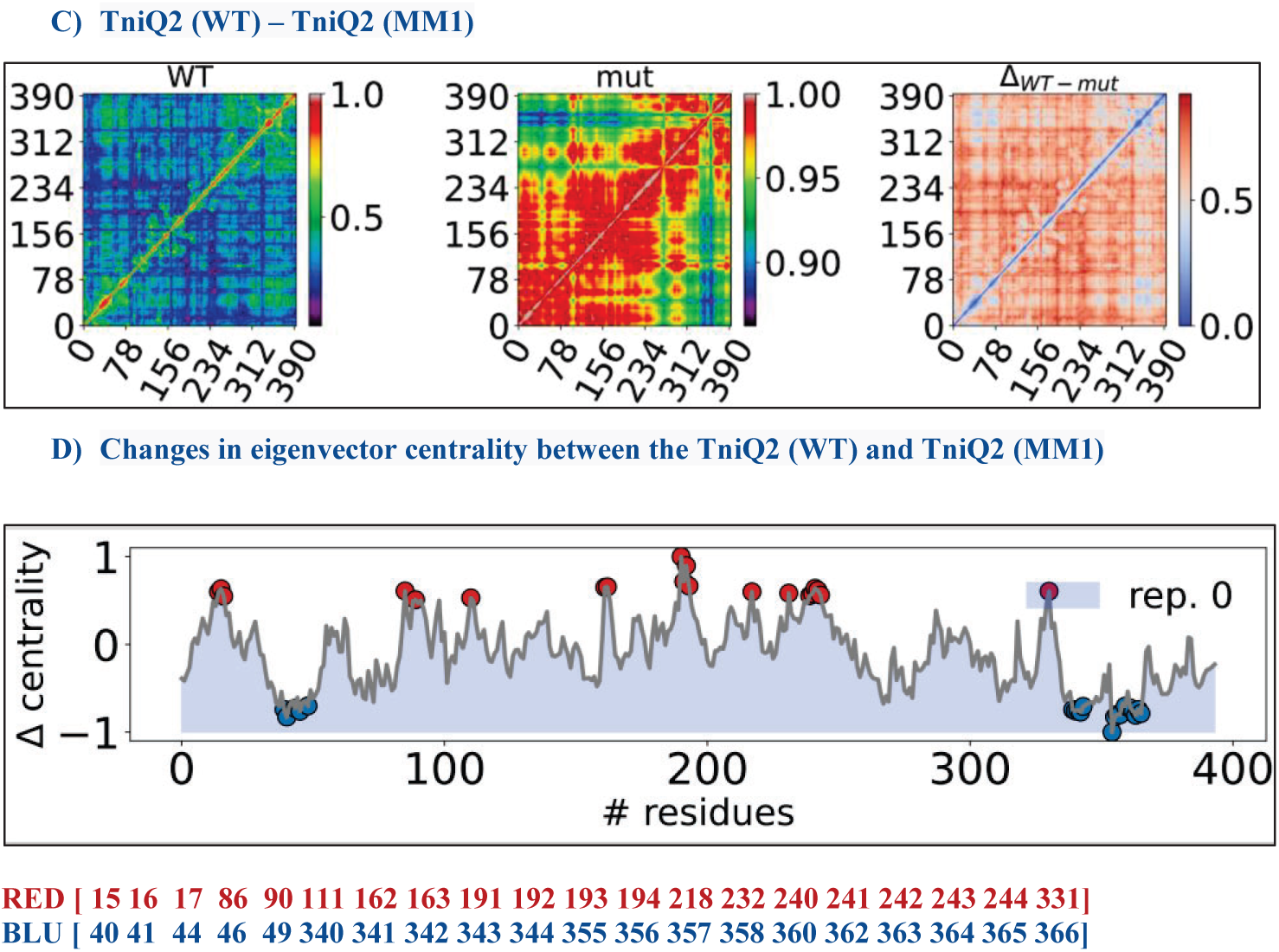
Correlation of TniQ1 and TniQ2 residues based on LMI metric for the full match model (WT), and the MM1 model and their differences.

**Fig.12.**
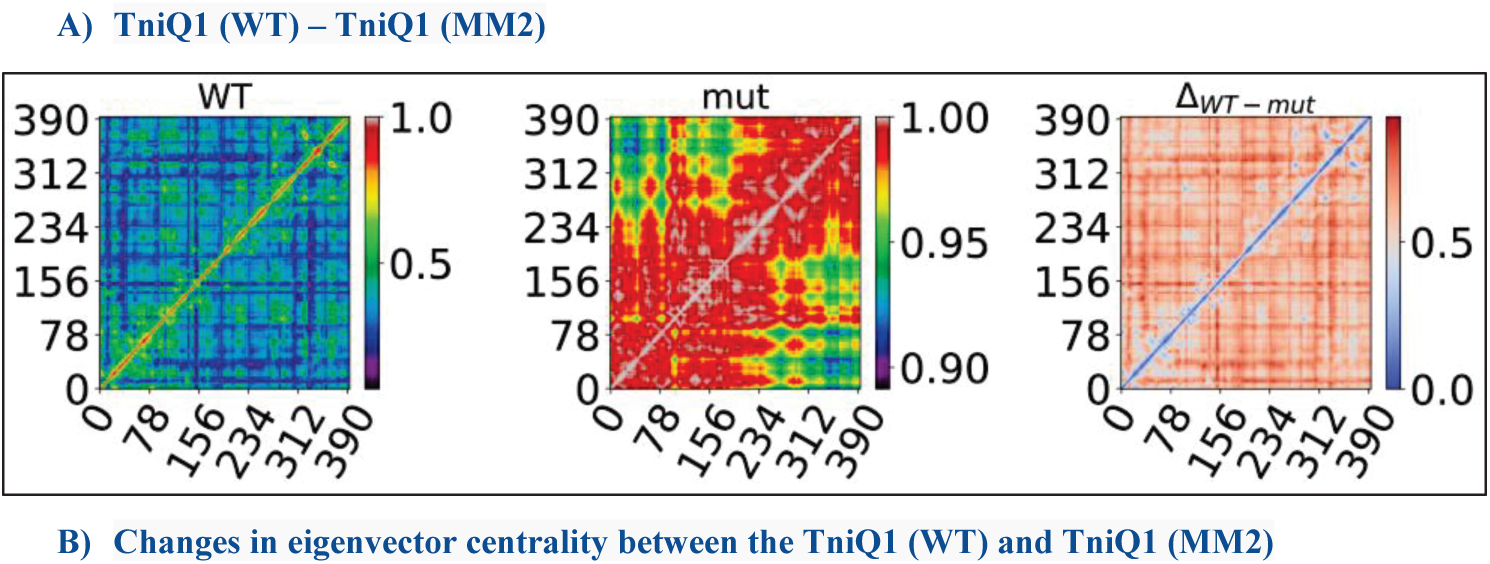
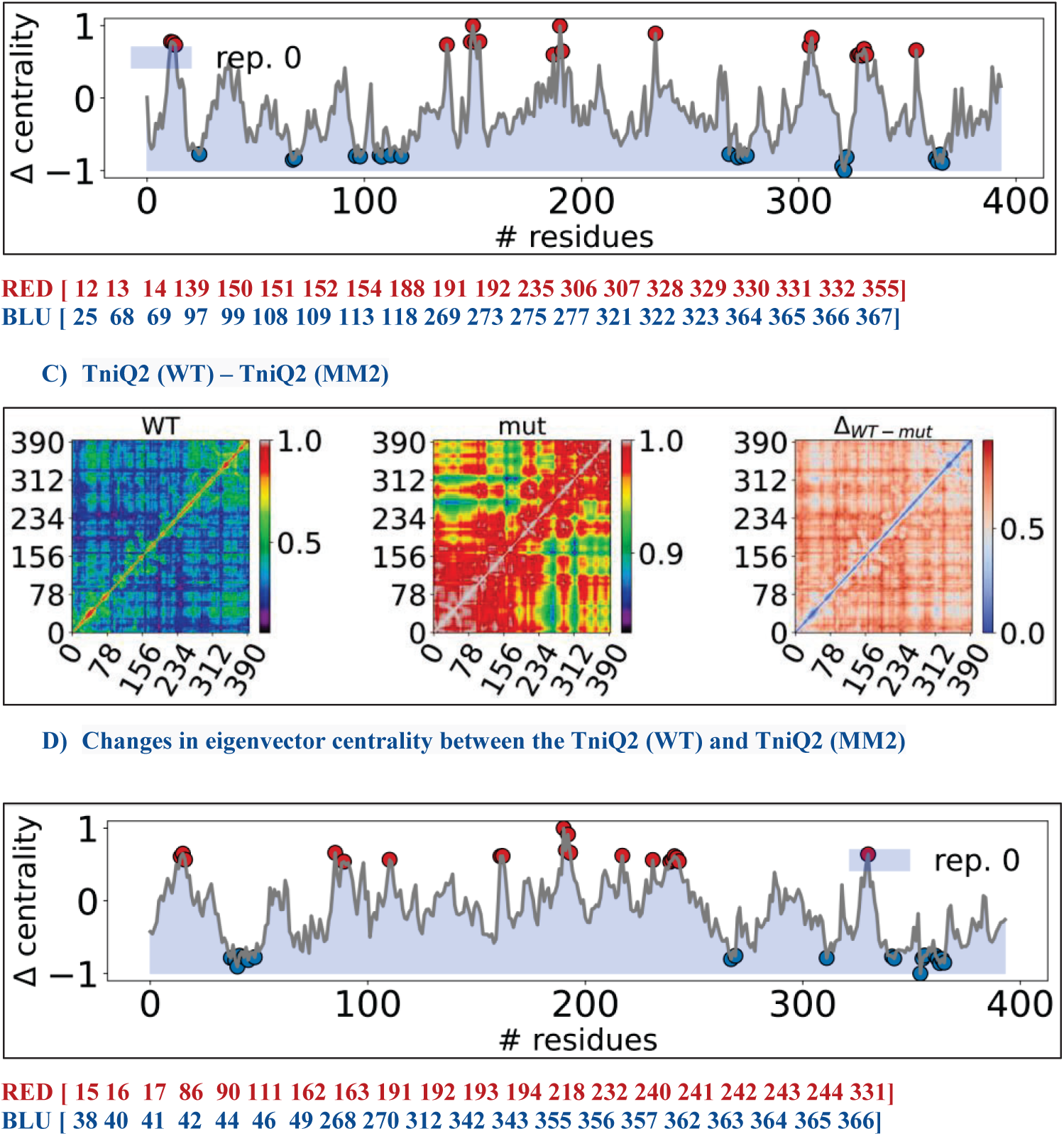
Correlation of TniQ1 and TniQ2 residues based on LMI metric for the full match model (WT), and the MM2 model and their differences.

**As shown in** **Fig.10**, the eigenvector centrality (EC) metric was utilized to identify key residues that played a significant role in the propagation of information throughout Cas8. The EC determines the importance of nodes within a network, based on the number and importance of their neighboring nodes, allowing for the identification of central residues with greater influence [29,31]. The introduction of a PAM-distal mismatch resulted in an increase in eigenvector centrality (EC) values for specific regions of Cas8, indicating that these regions became more important in the allosteric communications within Cas8. These regions were identified as the Cas8-NTD (amino acids 158-188) and the Cas8-CTD (amino acids 421-507), which were found to be particularly sensitive to the effects of the PAM-distal mismatch. Some residues experienced a decrease in the eigenvector centrality, which suggests that the node’s importance in the network has decreased. This could indicate that the node is no longer as critical to the communication pathways within the network or that its connections to other highly connected nodes have weakened.

The correlation matrices for the Cα-displacements from simulated trajectories of TniQ1 and TniQ2 in the wild-type (WT), MM1, MM2, and their differences with the changes in eigenvector centrality **are shown in** **Fig 11** **and** **12**

The red-circled residues have been identified as crucial for the TniQ dimer network, and their centrality undergoes significant increase upon exposure to PAM-distal mismatches. These residues may play a critical role in the interactions within the TniQ dimer network, and changes in their centrality values could reflect alterations in the network’s overall structure and function in response to the mismatches. In contrast, the blue-circled residues exhibit a decrease in eigenvector centrality values in the region. The change in the correlation patterns of the α-carbon displacements in the circled residues may have altered the importance of these residues in TniQ network.

To conduct further analysis, the MDigest tool was employed to calculate the correlation between the electrostatic potential across each frame in the trajectory of Cas8 and TniQ. [29]. To enable a direct comparison, we calculated the changes in electrostatic eigenvector centrality values (ΔEEC) induced by the mismatch, **as shown in Fig 13 and 14**

**Fig. 13.**
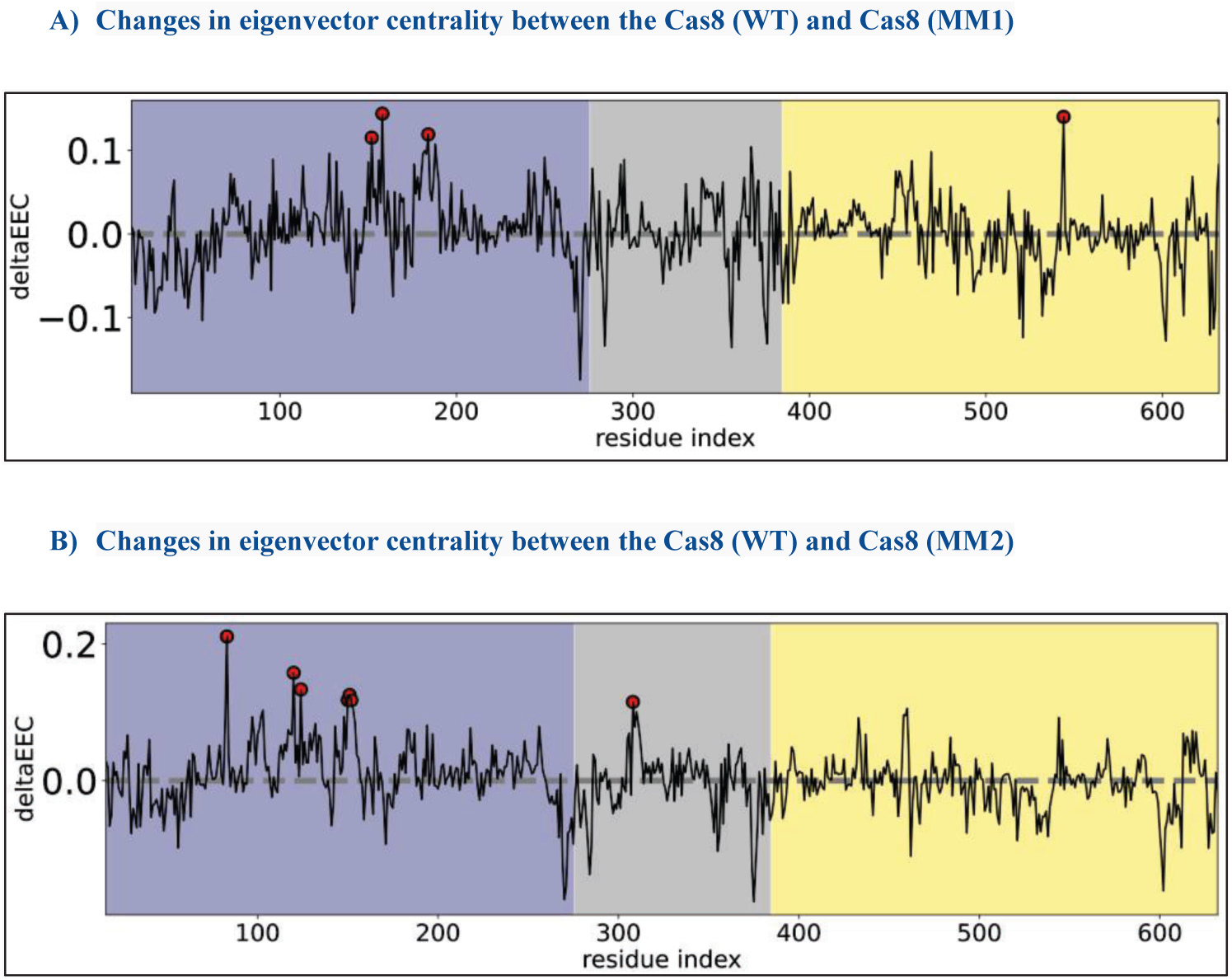
The change in electrostatic eigenvector centrality, as measured by the KS metric, between the Cas8 residues upon introducing mismatches within the DNA-RNA hybrid.

**Fig. 14.**
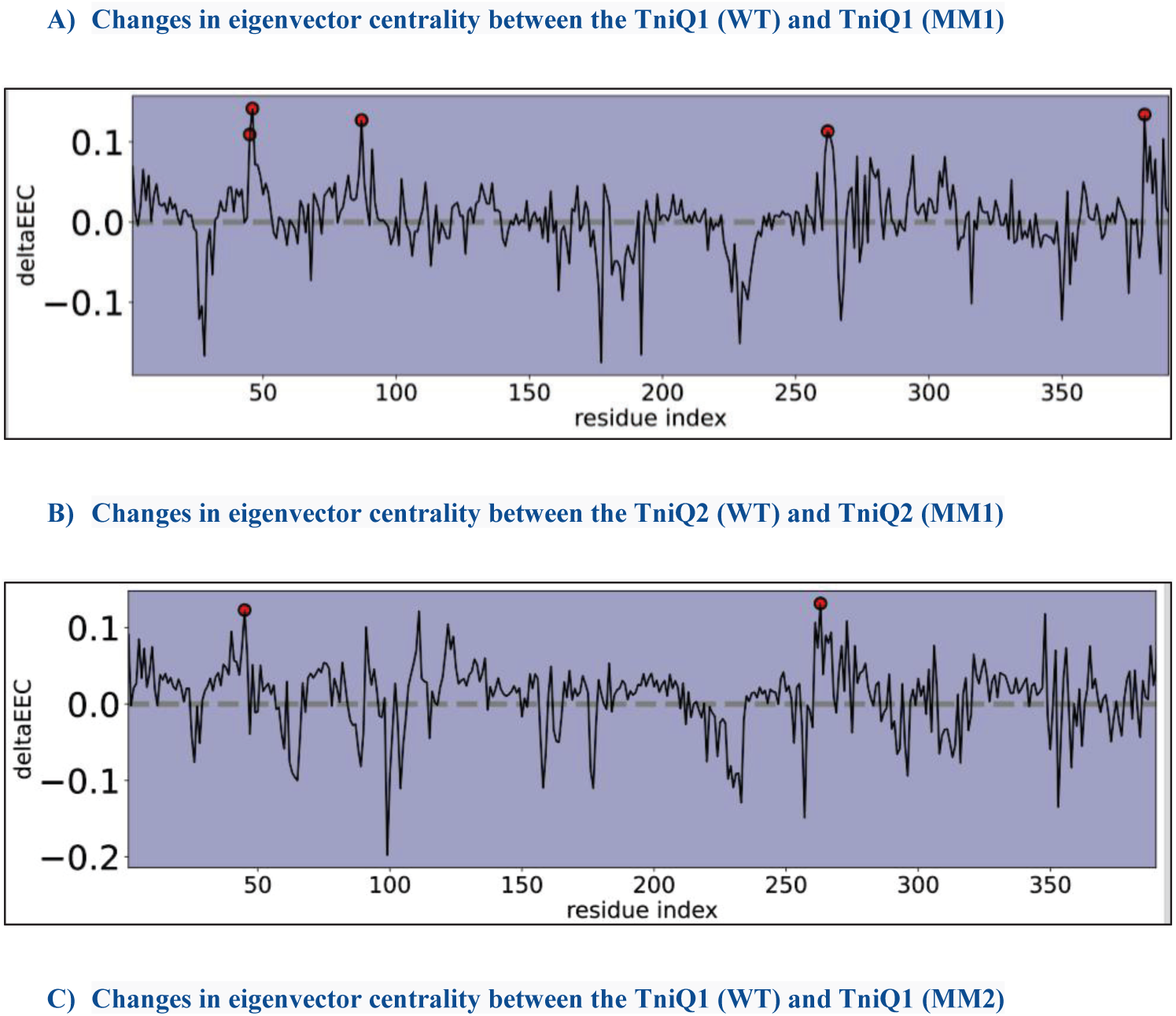
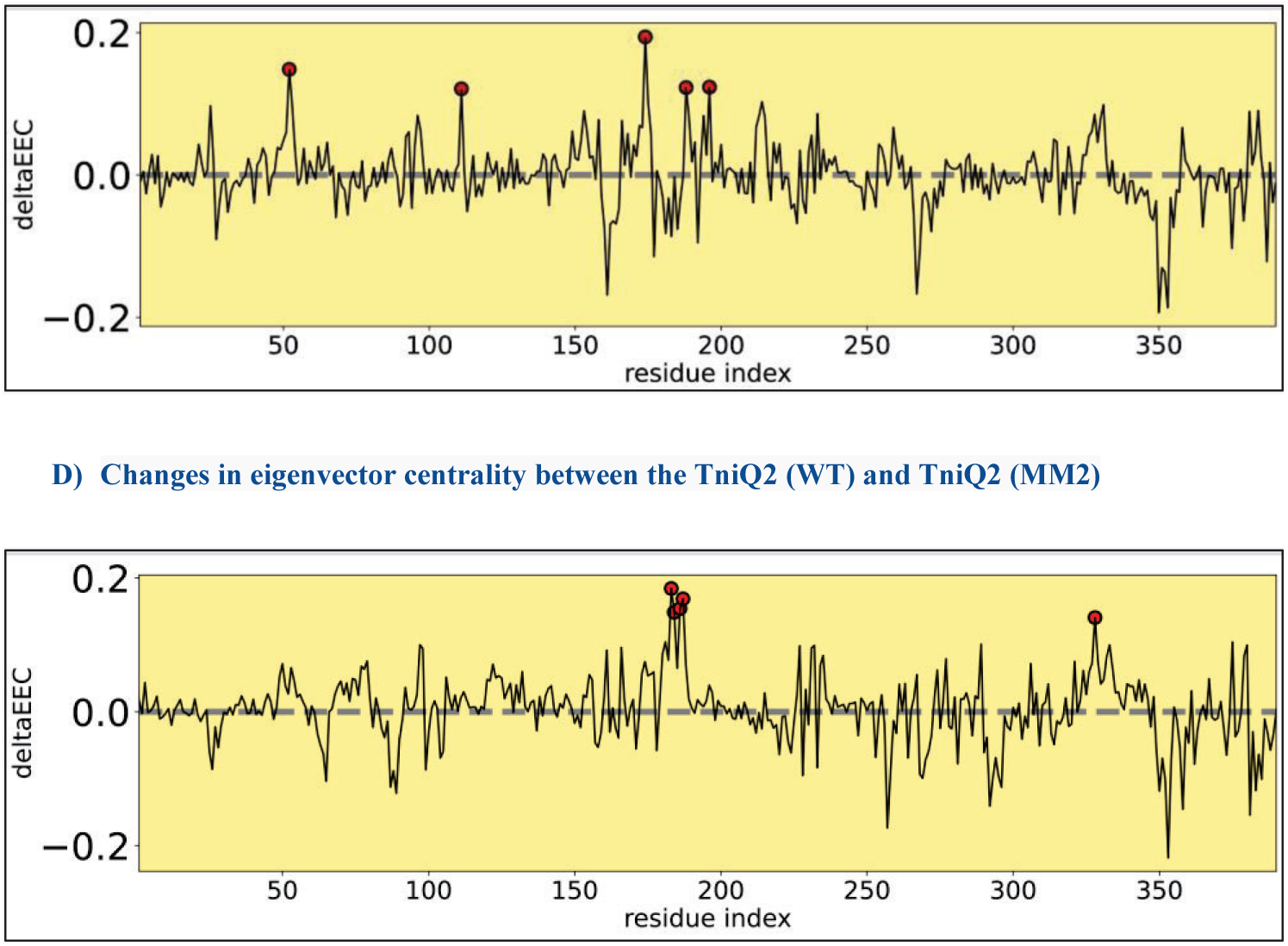
The change in electrostatic eigenvector centrality, as measured by the KS metric, between the TniQ residues in the full match and mismatch models.

Cas8 protein’s eigenvector centrality, as measured by LMI and KS metrics, indicates an overlap between amino acid residues 170, 177, 180, and 181. Therefore, these residues are likely to play a crucial role in allosteric communication within Cas8, as evidenced by significant changes in centrality under both mismatches.

This analysis revealed specific residues circled in red became significantly more central to the propagation of electrostatic information, which were affected by the PAM distal mismatch. However, the analysis did not show any overlap in the eigenvector centrality of the TniQ residues as measured by LMI and KS metrics, which are different measures of correlation that can complement each other. It is important to note that the LMI metric and the KS metric are both measures of pairwise interactions between residues in a protein, but they capture different aspects to describe the behavior of protein [29]. The LMI metric measures the linear mutual information between the α-carbon displacements of different residues, which can provide insight into the correlation patterns and communication pathways between different parts of the protein [28,29].

The KS metric, on the other hand, is based on the electrostatic interactions between residues and provides information about the stability and energetics of the protein structure [29,30]. Therefore, it is essential to consider multiple approaches and metrics when analyzing complex systems to fully understand their properties and behavior [29].

To gain further insight into the effect of the PAM distal mismatch on TniQ, we used the secondary structure community networks analysis to identify which regions were impacted by the mismatch. These secondary structure communities (nodes) were established by grouping adjacent residues that had the same secondary structure during most of the MD simulation. The edges between the nodes represent the interaction between the nodes and were defined using the KS-based GCC that was computed from KS energies throughout the trajectory, **as shown in Fig 15**

**Fig. 15.**
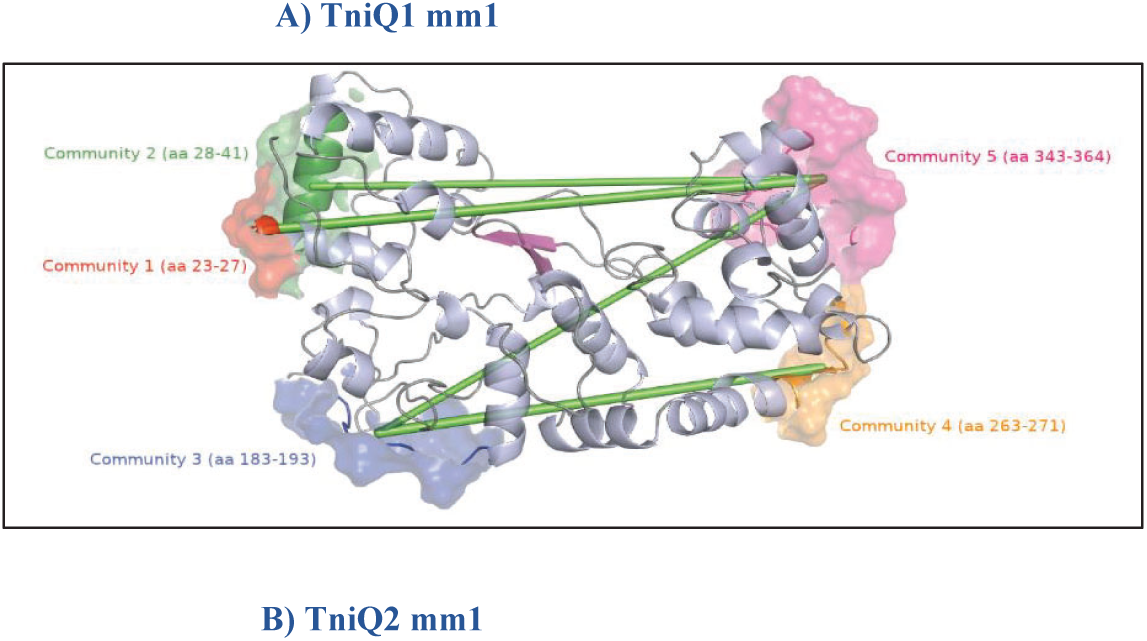
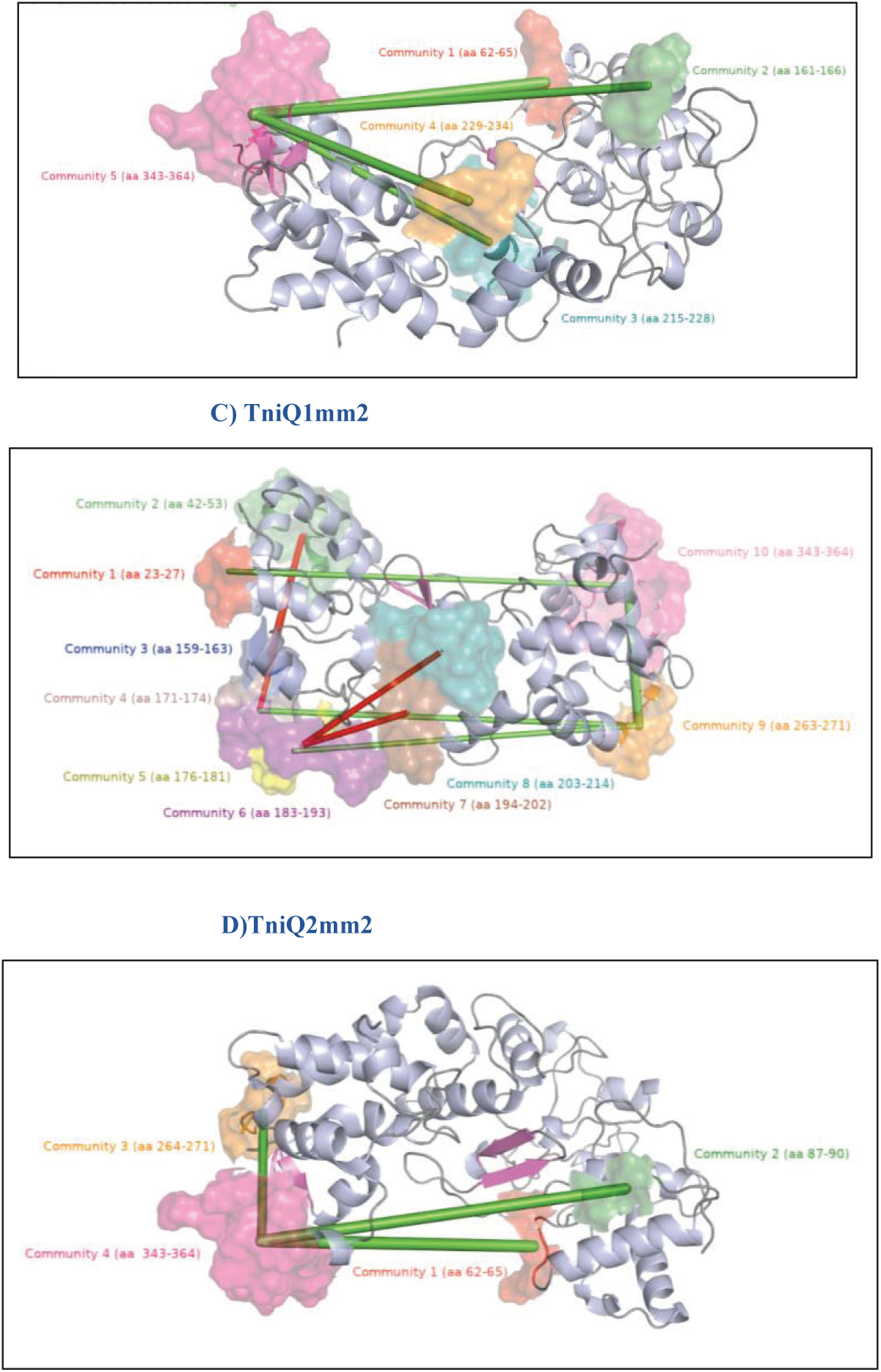
The secondary structure network of TniQ. The edges between the nodes are determined by changes in KS electrostatic energy caused by a PAM distal mismatch. Edges represent the difference in correlation coefficients between the full match model and mismatch models as the edges. Green edges indicate an increase in correlation, while red edges indicate a decrease in correlation.

When changes in the interactions between secondary structure communities are detected, the width of the edge connecting them is proportional to the magnitude of the change. If the change is positive, meaning there is an increase in the interaction strength between the communities, the edge is colored green. If the change is negative, meaning there is a decrease in the interaction strength between the communities, the edge is colored red.

The analysis revealed that introducing a PAM-distal mismatch at position 25-28 or (the MM1 model) resulted in Community 5, consisting of amino acids 343-364, being highly influential and central to the overall protein structure and function.

The PAM distal mismatch at position (29-32 or the MM2 model) revealed negative correlations between connected nodes in the network of TniQ1, as indicated by the presence of multiple edges with negative weights. For instance, the interaction between community 2 (amino acids 42-53) and community 4 (amino acids 171-174) weakens upon the mismatch. The absence of these two communities in the MM1 model may imply that their secondary structure is more unstable and prone to fluctuations, making it challenging to group the adjacent residues that share the same secondary structure. Interestingly, in both PAM-distal mismatches, the community of residues with amino acid numbers ranging from 343 to 364 appears to be highly connected to other communities in the protein structure, as indicated by its multiple edges to almost all nodes in the network. This suggests that this community is a key player in maintaining the overall stability and function of the protein, and any changes or disruptions to its structure or interactions could have significant impacts on the protein’s behavior.

The secondary structure community network can provide valuable insights into how the mismatch is affecting TniQ structure and dynamics, but further experimental and computational studies are necessary to determine the precise effects on protein function.

## Discussion

The RMSD analysis of the MD simulation trajectory provided insights into the conformational dynamics and stability of the of the INTEGRATE system. The MM1 model showed higher RMSD values for the backbone atoms compared to the other models, with Cas6 showing the highest fluctuations. Among the Cas7 subunits, Cas7.1, Cas7.2, and Cas7.5 exhibited the highest fluctuations in their RMSD values. Mismatches introduced with the DNA-RNA hybrid near Cas7.1 and Cas7.2 may have contributed to their behavior.

The results analyzed the impact of PAM-distal mismatches on the stability of the RNA-DNA hybrid in the CRISPR-Cas system. The analysis focused on the interactions between the protein subunits and the target strand of the DNA in the full match model and two PAM-distal mismatch models (MM1 model, where mismatches were introduced in positions 25-28, and MM2 model, where mismatches were introduced in positions 29-32). The results showed that some residues maintained their interactions with the DNA in all three models, while others experienced elongation or shortening of the distance between them and the DNA in the mismatch models. In some cases, new interactions were observed in the mismatch models. Interestingly, some salt bridge interactions were broken in the mismatch models while new ones were formed. The significance of the salt bridge formed by Lys230 in the full match and MM2 models lies in its potential role in stabilizing the RNA-DNA hybrid. In the full match model, the salt bridge was formed between Lys230 and the Guanine position at 64; however, this salt bridge was broken in both mismatch models. Interestingly, in the MM2 model, Lys230 formed a salt bridge with the Adenine positioned at 60 instead of the Guanine positioned at 64.

It was observed that in the MM1 model, the Thymine at position 58 was flipped. This finding suggests that spontaneous base flipping observed in the MM1 model could potentially be a mechanism for distinguishing between normal and damaged Watson-Crick base pairs [32].

In the full match and MM2 models, evidence supports that Cas8 helical bundle undergoes rotation and forms interactions with the TniQ dimer, which is subsequently stabilized by Cas7.2 interactions. This observation suggests that the stable rotation of Cas8-HB is an essential and conserved mechanism involved in the TniQ interaction and the DNA transposition process. However, the MM1 model does not show evidence of Cas8-HB rotation, which supports the suggestion that these positions are sensitive and essential for the proper functioning of the DNA transposition process, as the introduction of a mismatch at these positions can disrupt the interaction between Cas8-HB and TniQ.

Our results suggest that the presence of a PAM-distal mismatch can disrupt the electrostatic information network and alter the allosteric communication within Cas8 and TniQ dimers. By utilizing network analysis techniques such as eigenvector centrality and correlation analysis, we were able to identify specific amino acid residues that play a critical role in the allosteric communication mechanism. The community of residues ranging from 343 to 364 in the TniQ dimer appears to be highly connected to other communities, indicating that it may play a central role in the protein’s overall stability and function. This community could be i8nvolved in important interactions and structural features of the protein that contribute to its function. It should be noted that this particular region is a component of the alpha helix-turn-alpha helix (HTH) domain, which is anticipated to engage with the Cas8-HB [9].

## Conclusion

The INTEGRATE system is highly sensitive to mismatches in the PAM-proximal region of the crRNA, which significantly reduces transposition efficiency [7]. Mismatches in the PAM-distal region also affect the system, particularly at positions 25-28, which can impede the conformational changes necessary for TniQ to recruit TnsC [7].

The study used all-atom molecular dynamics simulations to investigate the impact of PAM-distal mismatches on the stability of the RNA-DNA hybrid in the CRISPR-Cas system. Three independent models were analyzed to provide a comprehensive view of the system’s conformational dynamics and stability. The results showed that mismatches in the PAM distal region can affect the stability of the RNA-DNA hybrid and disrupt the interactions between the protein subunits and the target strand of the DNA.

Our results revealed that the stable rotation of Cas8-HB is an essential mechanism in the TniQ interaction and the DNA transposition process. This observation is consistent with previous studies that have highlighted the importance of Cas8-HB rotation in type I CRISPR-Cas systems [19].

Overall, the network analysis techniques used in this study have provided valuable insights into the communication pathways and critical amino acid residues involved in the DNA transposition process in Cas8 and TniQ dimers. The results suggest that PAM-distal mismatches can affect the communication pathways and highlight the importance of specific amino acids in maintaining the stability and function of the protein structures. Furthermore, the highly connected community of residues ranging from 343 to 364 in TniQ may play a critical role in maintaining the overall stability and function of the protein. The identified amino acid residues and communication pathways within Cas8 and TniQ dimers provide valuable insights into the mechanisms involved in the DNA transposition process and how they are affected by PAM-distal mismatches.

However, further research is needed to fully understand the conformational changes required for R-loop locking and TniQ activation. The findings presented in this study will be useful for designing strategies to improve the efficiency and accuracy of the INTEGRATE system in genome engineering applications.

## Acknowledgement

Our research was made possible through the use of computational resources provided by the Shaheen supercomputer at King Abdullah University of Science and Technology (KAUST), and the XSEDE computational resources at Pittsburgh Supercomputing Center (PSC).

## References

1. Jansen R, Van Embden JDA, Gaastra W, Schouls LM. Identification of genes that are associated with DNA repeats in prokaryotes. Mol Microbiol 2002;43. 10.1046/j.1365-2958.2002.02839.x.

2. Mojica FJM, Díez-Villaseñor C, Soria E, Juez G. Biological significance of a family of regularly spaced repeats in the genomes of Archaea, Bacteria and mitochondria. Mol Microbiol 2000;36. 10.1046/j.1365-2958.2000.01838.x.

3. Makarova KS, Wolf YI, Koonin E V. Classification and Nomenclature of CRISPR-Cas Systems: Where from Here? Cris J 2018;1. 10.1089/crispr.2018.0033.

4. Fu Y, Foden JA, Khayter C, Maeder ML, Reyon D, Joung JK, et al. High-frequency off-target mutagenesis induced by CRISPR-Cas nucleases in human cells. Nat Biotechnol 2013;31. 10.1038/nbt.2623.

5. Klompe SE, Sternberg SH. Harnessing “A Billion Years of Experimentation”: The Ongoing Exploration and Exploitation of CRISPR–Cas Immune Systems. Cris J 2018;1:141–58. 10.1089/crispr.2018.0012.

6. Peters JE, Makarova KS, Shmakov S, Koonin E V. Recruitment of CRISPR-Cas systems by Tn7-like transposons. Proc Natl Acad Sci U S A 2017;114:E7358–66. 10.1073/pnas.1709035114.

7. Klompe SE, Vo PLH, Halpin-Healy TS, Sternberg SH. Transposon-encoded CRISPR–Cas systems direct RNA-guided DNA integration. Nature 2019;571:219–25. 10.1038/s41586-019-1323-z.

8. Shen Y, Gomez-Blanco J, Petassi MT, Peters JE, Ortega J, Guarné A. Structural basis for DNA targeting by the Tn7 transposon. Nat Struct Mol Biol 2022;29:143–51. 10.1038/s41594-022-00724-8.

9. Li Z, Zhang H, Xiao R, Chang L. Cryo-EM structure of a type I-F CRISPR RNA guided surveillance complex bound to transposition protein TniQ. Cell Res 2020;30:179–81. 10.1038/s41422-019-0268-y.

10. van der Oost J, Mougiakos I. First structural insights into CRISPR-Cas-guided DNA transposition. Cell Res 2020;30:193–4. 10.1038/s41422-020-0284-y.

11. Jia N, Xie W, de la Cruz MJ, Eng ET, Patel DJ. Structure–function insights into the initial step of DNA integration by a CRISPR–Cas–Transposon complex. Cell Res 2020;30:182–4. 10.1038/s41422-019-0272-2.

12. Halpin-Healy TS, Klompe SE, Sternberg SH, Fernández IS. Structural basis of DNA targeting by a transposon-encoded CRISPR–Cas system. Nature 2020;577:271–4. 10.1038/s41586-019-1849-0.

13. Wang B, Xu W, Yang H. Structural basis of a Tn7-like transposase recruitment and DNA loading to CRISPR-Cas surveillance complex. Cell Res 2020;30:185–7. 10.1038/s41422-020-0274-0.

14. Songailiene I, Rutkauskas M, Sinkunas T, Manakova E, Wittig S, Schmidt C, et al. Decision-Making in Cascade Complexes Harboring crRNAs of Altered Length. Cell Rep 2019;28. 10.1016/j.celrep.2019.08.033.

15. Van Erp PBG, Jackson RN, Carter J, Golden SM, Bailey S, Wiedenheft B. Mechanism of CRISPR-RNA guided recognition of DNA targets in Escherichia coli. Nucleic Acids Res 2015;43. 10.1093/nar/gkv793.

16. Zheng Y, Li J, Wang B, Han J, Hao Y, Wang S, et al. Endogenous Type I CRISPR-Cas: From Foreign DNA Defense to Prokaryotic Engineering. Front Bioeng Biotechnol 2020;8:1–17. 10.3389/fbioe.2020.00062.

17. Hille F, Richter H, Wong SP, Bratovič M, Ressel S, Charpentier E. The Biology of CRISPR-Cas: Backward and Forward. Cell 2018;172:1239–59. 10.1016/j.cell.2017.11.032.

18. Xue C, Sashital DG. Mechanisms of Type I-E and I-F CRISPR-Cas Systems in Enterobacteriaceae. EcoSal Plus 2019;8. 10.1128/ecosalplus.esp-0008-2018.

19. Rollins MCF, Chowdhury S, Carter J, Golden SM, Miettinen HM, Santiago-Frangos A, et al. Structure Reveals a Mechanism of CRISPR-RNA-Guided Nuclease Recruitment and Anti-CRISPR Viral Mimicry. Mol Cell 2019;74. 10.1016/j.molcel.2019.02.001.

20. Jo S, Kim T, Iyer VG, Im W. CHARMM-GUI: A web-based graphical user interface for CHARMM. J Comput Chem 2008;29. 10.1002/jcc.20945.

21. Lu XJ, Olson WK. 3DNA: A software package for the analysis, rebuilding and visualization of three-dimensional nucleic acid structures. Nucleic Acids Res 2003;31:5108–21. 10.1093/nar/gkg680.

22. Van Der Spoel D, Lindahl E, Hess B, Groenhof G, Mark AE, Berendsen HJC. GROMACS: Fast, flexible, and free. J Comput Chem 2005;26. 10.1002/jcc.20291.

23. Vangone A, Spinelli R, Scarano V, Cavallo L, Oliva R. COCOMAPS: A web application to analyze and visualize contacts at the interface of biomolecular complexes. Bioinformatics 2011;27:2915–6. 10.1093/bioinformatics/btr484.

24. Xiao Y, Luo M, Hayes RP, Kim J, Ng S, Ding F, et al. Structure Basis for Directional R-loop Formation and Substrate Handover Mechanisms in Type I CRISPR-Cas System. Cell 2017;170. 10.1016/j.cell.2017.06.012.

25. Bosshard HR, Marti DN, Jelesarov I. Protein stabilization by salt bridges: Concepts, experimental approaches and clarification of some misunderstandings. J Mol Recognit 2004;17. 10.1002/jmr.657.

26. Shields R. Cultural Topology: The Seven Bridges of Königsburg, 1736. Theory, Cult Soc 2012;29. 10.1177/0263276412451161.

27. Bahar I, Atilgan AR, Erman B. Direct evaluation of thermal fluctuations in proteins using a single-parameter harmonic potential. Fold Des 1997;2. 10.1016/S1359-0278(97)00024-2.

28. Lange OF, Grubmüller H. Generalized correlation for biomolecular dynamics. Proteins Struct Funct Genet 2006;62. 10.1002/prot.20784.

29. Federica Maschietto BAGKVSB. MDiGest: A Python Package for Describing Allostery from Molecular Dynamics Simulations n.d.

30. Kabsch W, Sander C. Dictionary of protein secondary structure: Pattern recognition of hydrogen-bonded and geometrical features. Biopolymers 1983;22. 10.1002/bip.360221211.

31. Maschietto F, Zavala E, Allen B, Loria JP, Batista V. MptpA Kinetics Enhanced by Allosteric Control of an Active Conformation. J Mol Biol 2022;434. 10.1016/j.jmb.2022.167540.

32. Yin Y, Yang L, Zheng G, Gu C, Yi C, He C, et al. Dynamics of spontaneous flipping of a mismatched base in DNA duplex. Proc Natl Acad Sci U S A 2014;111. 10.1073/pnas.1400667111.

